# A programmable probiotic encapsulation system enhances therapeutic delivery *in vivo*

**DOI:** 10.1101/2021.08.03.454638

**Authors:** Tetsuhiro Harimoto, Jaeseung Hahn, Yu-Yu Chen, Jongwon Im, Joanna Zhang, Nicholas Hou, Fangda Li, Courtney Coker, Kelsey Gray, Nicole Harr, Sreyan Chowdhury, Kelly Pu, Clare Nimura, Nicholas Arpaia, Kam Leong, Tal Danino

**Affiliations:** Department of Biomedical Engineering, Columbia University, New York, NY 10027, USA; Department of Microbiology and Immunology, Vagelos College of Physicians and Surgeons, Columbia University, New York, NY 10032, USA; Herbert Irving Comprehensive Cancer Center, Columbia University, New York, NY 10032, USA; Department of Systems Biology, Columbia University Medical Center, New York, NY 10032, USA; Data Science Institute, Columbia University, New York, NY 10027, USA

## Abstract

Recent advances in therapeutic modulation of human microbiota have driven new efforts to engineer living microbial medicines using synthetic biology. However, a long-standing challenge for live bacterial therapies is balancing the high dose required to achieve robust efficacy with the potential for sepsis. Here, we developed a genetically encoded microbial encapsulation system with tunable and dynamic expression of surface capsular polysaccharides to enhance therapeutic delivery. Following a synthetic small RNA knockdown screen of the capsular biosynthesis pathway, we constructed synthetic gene circuits that regulate bacterial encapsulation based on sensing the levels of environmental inducer, bacterial density, and blood pH. The induced encapsulation system enabled tunable immunogenicity and survivability of the probiotic *Escherichia coli*, resulting in increased maximum tolerated dose and enhanced efficacy in murine cancer models. Furthermore, triggering *in situ* encapsulation was found to increase microbial translocation between mouse tumors, leading to efficacy in distal tumors. The programmable encapsulation system demonstrates a new approach to control microbial therapeutic profiles *in vivo* using synthetic biology.

## Main Text

The microbiome plays numerous functional roles in human health and subsequently has led to focused interest in the use of live bacteria to treat disease^1,2^. Since microbes can be engineered as intelligent living medicines that sense and respond to environments, they can colonize niches in the gastrointestinal tract^3,4^, mouth^5^, skin^6^, lung^7^, and tumors^8,9^, and locally deliver therapeutics. However, host toxicity from live bacteria has been shown to limit tolerated dose and efficacy, in some cases leading to termination of clinical trials^10–13^. Moreover, unlike conventional drug carriers, the unique abilities of bacteria to continuously proliferate, chemotax, and produce therapeutic payloads in disease sites necessitates robust and temporal control of bacterial pharmacokinetics *in vivo*. One approach to circumvent toxicity is the generation of genetic knockouts of immunogenic bacterial surface antigens such as lipopolysaccharide (LPS), but this strategy can result in permanent strain attenuation and reduced colonization, as seen in clinical trials of bacteria cancer therapy^11,14,15^. Surface modulation has been widely utilized in cloaking drug delivery vehicles^16^, and thus an alternative strategy is the synthetic coating of microbial surfaces with molecules such as alginate^17,18^, chitosan^17^, polydopamine^19^, lipids^20–22^, and nanoparticles^23^. However, these one-time, static modifications of bacteria do not allow for *in situ* modulation and can lead to uncontrolled growth, off-target toxicity, or compromised cellular function resulting in reduced efficacy.

Here, we present a tunable microbial surface engineering strategy using synthetic gene circuits to dynamically control bacterial interactions with their surrounding environment. We focused on bacterial surface capsular polysaccharides (CAP), a natural extracellular biopolymer that coats the extracellular membrane and protects microbes from a variety of environmental conditions^24^. By applying a synthetic biology approach to engineer CAP biosynthesis, we constructed programmable CAP expression systems that sense environmental cues and controllably modulate the bacterial surface, thereby modulating bacterial interaction with antimicrobials, bacteriophage, acidity, and host immunity. This design allows precise control over bacterial immunogenicity and survivability *in vivo*, enabling novel drug delivery strategies such as enhanced dosing and *in situ* trafficking to maximize therapeutic efficacy and safety (Fig. 1).

**Figure 1:**
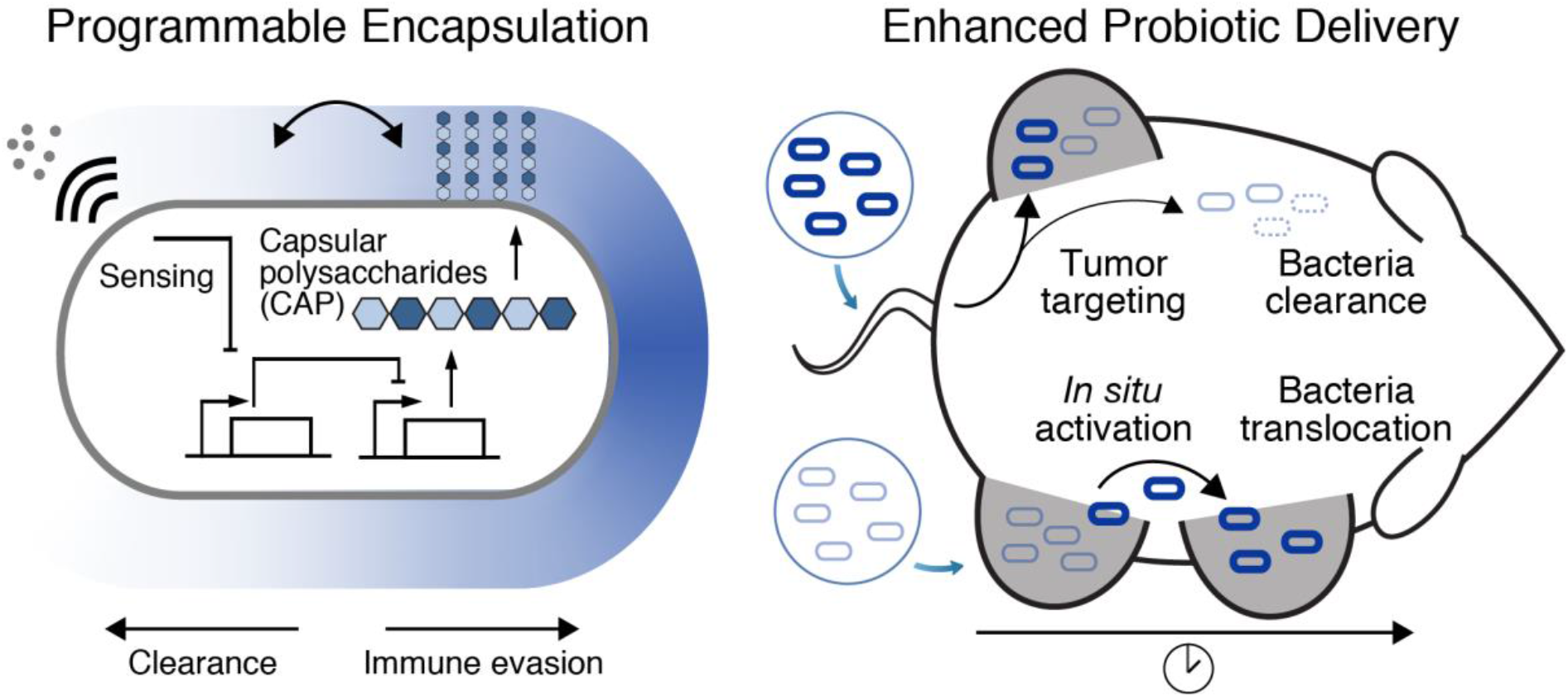
Programmable capsular polysaccharides (CAP) system for control over bacterial encapsulation and *in vivo* delivery profiles. We engineered the biosynthetic pathway of bacterial CAP for tunable and dynamic surface modulation of the probiotic *E. coli* Nissle 1917 with synthetic gene circuits. The CAP system modulates bacterial immunogenicity and survivability *in vivo*. By balancing these factors, the programmable CAP system is capable of reducing toxicity related to systemic bacterial administration and enables inducible bacterial translocation between tumors.

### sRNA knockdown screen identifies key regulators of CAP synthesis

Since various bacteria have been utilized for therapeutic applications, we compared immunogenicity and viability of several *E. coli* and *S. typhimurium* strains. Here *E. coli* Nissle 1917 (EcN), a probiotic strain with favorable clinical profiles^25^, demonstrated high viability in human whole blood with minimal cytokine induction (Supplementary Fig. 1a,b). Because the K5-type CAP of EcN has been shown to alter interaction with host immune systems^26–29^, we chose to genetically modify its biosynthetic pathway^30,31^. K5-type CAP produced from EcN, also known as heparosan, is composed of a polymer chain of alternating β-D-glucuronic acid (GlcA) and N-acetyl-α-D-glucosamine (GlcNAc), attached to 3-deoxy-D-manno-oct-2-ulosonic acid (Kdo) linker (Fig. 2a). Glycotransferases of *kfiABCD* genes polymerize alternating GlcA and GlcNAc subunits. *kpsCSFU* genes are responsible for synthesis of the poly-Kdo linker on the terminal lipid, and CAP is transported to the cellular surface by *kpsEDMT* genes. While individual functions of the CAP genes have been investigated, engineering tunable and dynamic control of this system remains unexplored.

**Figure 2:**
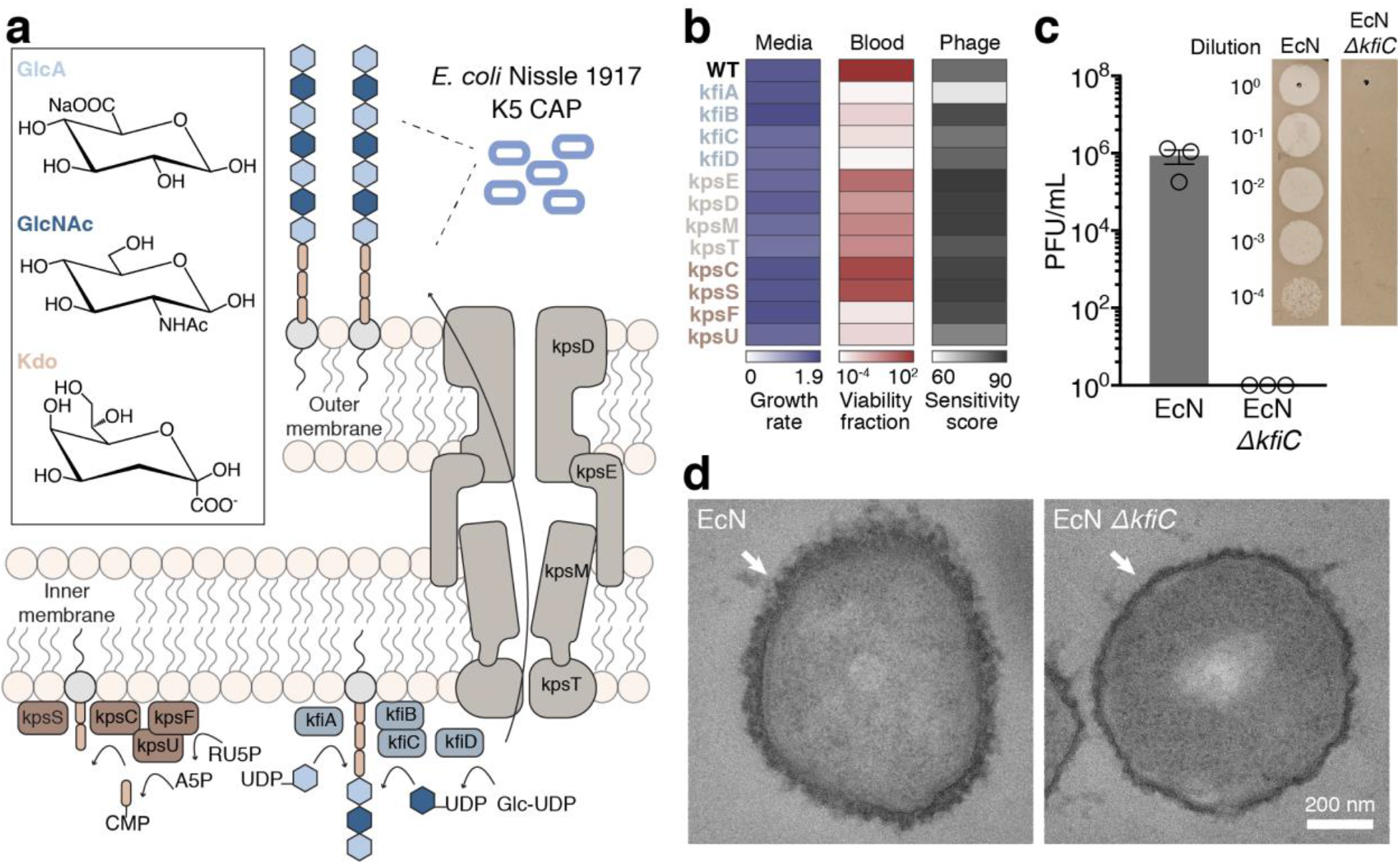
sRNA knockdown screen identifies key genes in capsular polysaccharides (CAP) biosynthesis. **a,** Schematics of K5 CAP biosynthesis in EcN. CAP is composed of an alternating polymer chain of GlcA and GlcNAc connected to a poly-KDO linker. Subsequently CAP is transported from the inner bacteria membrane to the outer membrane. **b,** Quantification of microbial growth parameters of EcN knockdown (KD) strains in nutrient, blood, or phage containing media. Growth rate denotes maximum specific growth rate (hour^-1^) obtained by fitting growth curve to measured OD600 over time. Blood viability is defined as fraction of bacterial CFU after 6 hours incubation in human blood over inoculated bacterial CFU. Phage sensitivity is calculated by area under the curve of bacterial turbidity over 6 hours of incubation with LB media containing ΦK1-5. **c,** Phage sensitivity of EcN and EcN Δ*kfiC*. Plaque forming assay demonstrates complete absence of infection and lysis in EcN Δ*kfiC*. The representative images show difference between serially-diluted plaque forming units (PFU) of bacteria with and without CAP. **d,** TEM images showing CAP encapsulation of the cellular outer surface. *kfiC* knockout results in the absence of CAP nanostructure on the cell surface of EcN Δ*kfiC*. White arrows indicate cell surface.

We sought to identify key CAP genes capable of altering response to antibacterial factors encountered during therapeutic delivery. To do so, we generated a library of knockdown (KD) strains using synthetic small RNAs (sRNAs) that reduce expression of *kfi* and *kps* genes via complementary binding to mRNAs^32^. To initially assess the impact of downregulating each gene, we screened the growth of KD strains in (1) nutrient-rich media, (2) human whole blood, and (3) CAP-targeting phage. Growth in nutrient-rich media showed little variation in maximum specific growth rates (μm) from the wild-type EcN strain (expressing CAP) (Fig. 2b, Supplementary Fig. 2a), suggesting that the downregulation of the targeted genes in the CAP biosynthetic pathway does not greatly affect the fitness of EcN in the absence of environmental threats. However, we observed significantly reduced viability of KD strains compared to EcN after incubation in whole blood for 0.5 hours (Supplementary Fig. 2b). After a 6-hour incubation in whole blood, KD strains in CAP synthesis (*kfi* genes and *kpsFU*) exhibited lower viability compared to KD strains in CAP transport (*kpsEDMT*) (Fig. 2b). To assess whether each gene KD causes a complete or partial loss of CAP in KD strains, we used lytic bacteriophage ΦK1-5 that specifically binds to heparosan of EcN^29,33^. Bacteria that express residual levels of heparosan CAP are susceptible to this phage, but complete loss of CAP confers immunity. We quantified phage sensitivity of each strain by measuring area under the curve of the phage-inoculated growth curve^34^ and observed that sRNA KD of most CAP genes did not alter phage sensitivity compared to control EcN (Fig. 2b, Supplementary Fig. 2c), suggesting some level of CAP is still present in KD strains. To abrogate the effect of residual CAP gene expression, we next constructed a library of knockout (KO) strains by deleting CAP synthesis genes from EcN genome using Lambda Red recombineering system. All *kfi* KO strains resulted in complete phage immunity (Supplementary Fig. 2d), supporting loss of CAP expression. Importantly, KO strains demonstrated significantly increased blood sensitivity (Supplementary Fig. 2e), indicating that CAP levels can be genetically tuned to alter sensitivities to antibacterial factors.

On the basis of the above results, we chose to further characterize *kfiC*, a well-studied gene that encodes an essential glycotransferase of GlcA^26,29^. Downregulation of *kfiC* via sRNA KD sensitized bacteria in blood, suggesting its key role in regulating bacterial protection. Deletion of *kfiC* resulted in the highest enhancement in blood sensitivity, indicating that the level of protection can be altered by controlling gene expression. To confirm loss of CAP from the bacterial surface, we characterized surface properties of EcN Δ*kfiC* strain. Phage plaque formation assay confirmed complete immunity against ΦK1-5 (Fig. 2c). We also purified and detected bacterial polysaccharides using sodium dodecyl sulfate–polyacrylamide gel electrophoresis (SDS-PAGE) followed by CAP staining with alcian blue. Compared to EcN that produced strong staining with alcian blue at ~180 kDa, EcN Δ*kfiC* produced no visible band (Supplementary Fig. 3a). We next characterized the morphological changes in bacterial surface using transmission electron microscopy (TEM) with ruthenium red staining. CAP was visible as an ~80 nm thick layer of polysaccharides coating outside of the cellular membrane. In contrast, EcN Δ*kfiC* had diminished size of the polysaccharides layer at ~40 nm (Fig. 2d, Supplementary Fig. 3a,b). We then investigated capability of CAP to protect cells from wide range of antimicrobial factors. In addition to the modified sensitivity to human whole blood and bacteriophage, EcN Δ*kfiC* demonstrated a significant reduction in cellular protection against panels of antibiotics (spectinomycin, ampicillin, gentamicin, kanamycin, streptomycin) and extreme acids (pH 2.5) compared to EcN (Supplementary Fig. 4a-i). Finally, we evaluated general applicability of the approach in other CAP systems. We deleted homologous genes in different *E. coli* strains expressing K1 and K5 CAP (*neuC* and *kfiC*, respectively) and showed alteration in environmental sensitivity (Supplementary Fig. 5a-c). Together, these results demonstrate that loss of CAP modifies cellular surface structure and protection against antimicrobial factors.

### Construction of tunable and reversible programmable CAP

We next constructed a programmable CAP system that can sense and respond to induction stimuli and modulate cell surface properties. We cloned *kfiC* under the control of the *tac* promoter, which can be activated with the small-molecule inducer isopropyl-b-D-thiogalactopyranoside (IPTG) (Fig. 3a). Since the EcN genome encodes for constitutive *lacI* expression, we built a small library of plasmids with various copy numbers of *kfiC* to optimize for tight regulation of CAP production. EcN Δ*kfiC* transformed with the low (sc101 origin) copy number plasmid exhibited complete immunity against ΦK1-5 (Supplementary Fig. 6a), indicating tight repression at the basal level. Induction with IPTG rescued the phage sensitivity (Supplementary Fig. 6b), confirming inducible modulation of CAP on cellular surface.

**Figure 3:**
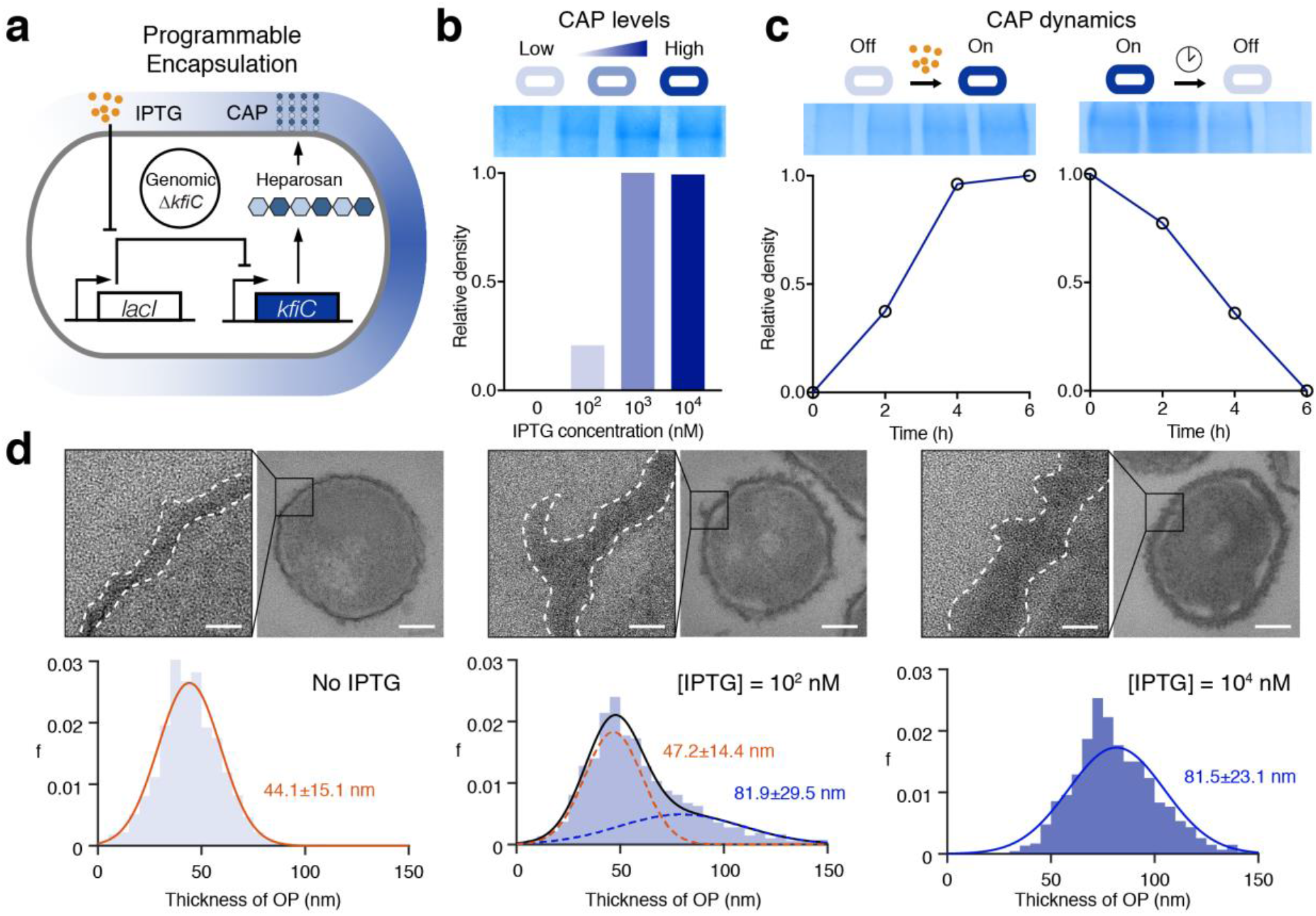
Design and characterization of the inducible capsular polysaccharides (iCAP) system. **a,** Inducible gene circuit diagram whereby the *kfiC* gene was cloned under the control of a *lac* promoter to allow inducible CAP expression via the small molecule IPTG. Copy number of the *kfiC* gene was modified to minimize basal *kfiC* expression. **b,** SDS-PAGE gel stained with Alcian blue showed elevating levels of CAP production corresponding to the IPTG concentration (top). The densitometric analysis of CAP bands demonstrated that CAP production reaches maximum at approximately 1 μM IPTG (bottom). **c,** SDS-page gels and densitometric analysis show CAP kinetics upon induction (left) and decay (right). **d,** Ruthenium red-stained TEM images showing change in CAP in titrating IPTG concentration. Histograms reveal shift in cellular outer layer thickness as IPTG concentration increases. Insets show representative images of bacteria and zoomed outer surface structure. Dotted lines indicate inner (white) and outer (gray) perimeters of CAP. Scale bar is 40 nm (left) or 200 nm (right) in each inset.

We examined tunability of this inducible CAP (iCAP) system by characterizing multiple induction conditions. SDS-PAGE showed increase in CAP production from EcN carrying the iCAP system (EcN iCAP) when incubated with elevating levels of IPTG (Fig. 3b). Co-incubation with ΦK1-5 also showed decreasing viability of EcN iCAP with elevating levels of IPTG (Supplementary Fig. 6c). We next used TEM to investigate the effect of the iCAP system on cell surface morphology (Fig. 3d). Increasing levels of IPTG shifted the mean bacterial membrane thickness from 44 nm to 81 nm, confirming tunable capability of the system. Intermediate iCAP activation at 100 nM revealed a bimodal distribution of the membrane thickness, suggesting that *kfiC* regulates the production level but not the length of the polysaccharide polymers. This result agreed with SDS-PAGE data that showed no difference in migration of CAP band depending on IPTG concentration (Fig. 3b).

We subsequently evaluated the dynamics of production and recovery of the iCAP system using a similar approach. Upon addition of IPTG, elevated CAP production was observed over time on SDS-PAGE, reaching near-maximum levels by 4 hours (Fig. 3c). Similarly, removal of IPTG resulted in gradual decrease in CAP until complete repression by 6 hours. We also tested iCAP dynamics via co-incubation with ΦK1-5. While uninduced EcN iCAP grew, induction with IPTG at the start of co-incubation resulted in a rapid lysis event at 3.5 hours (Supplementary Fig. 6d), demonstrating delayed CAP production with similar kinetics observed in SDS-PAGE. Collectively, these data highlight the programmable capability of CAP modulation on the bacterial surface.

### Programmable protection from host immunity

To build towards utilization of the programmable CAP system for therapeutic applications *in vivo*, we first tested the ability to exogenously control bacterial viability in human whole blood containing functional host bactericidal factors *in vitro*. Upon IPTG induction, we observed increased EcN iCAP survival compared to non-induced control (Fig. 4a). Increasing IPTG levels improved bacterial survival over a range of at least ~10^5^ fold, highlighting the tunable capability of the system. Since iCAP deactivation was observed after removing the inducer, we tracked bacterial survival over time after transiently activating EcN iCAP at varying IPTG concentration. We were able to modulate the rate of bacterial clearance from blood by titrating levels of IPTG (Fig. 4b). Wildtype EcN persisted for >6 hours, while EcN Δ*kfiC* quickly decreased to the levels under the limit of detection (LOD ~10^2^ CFU/mL) within the first 0.5 hour. A protective role of CAP in mouse whole blood was also observed (Supplementary Fig. 7a,b).

**Figure 4:**
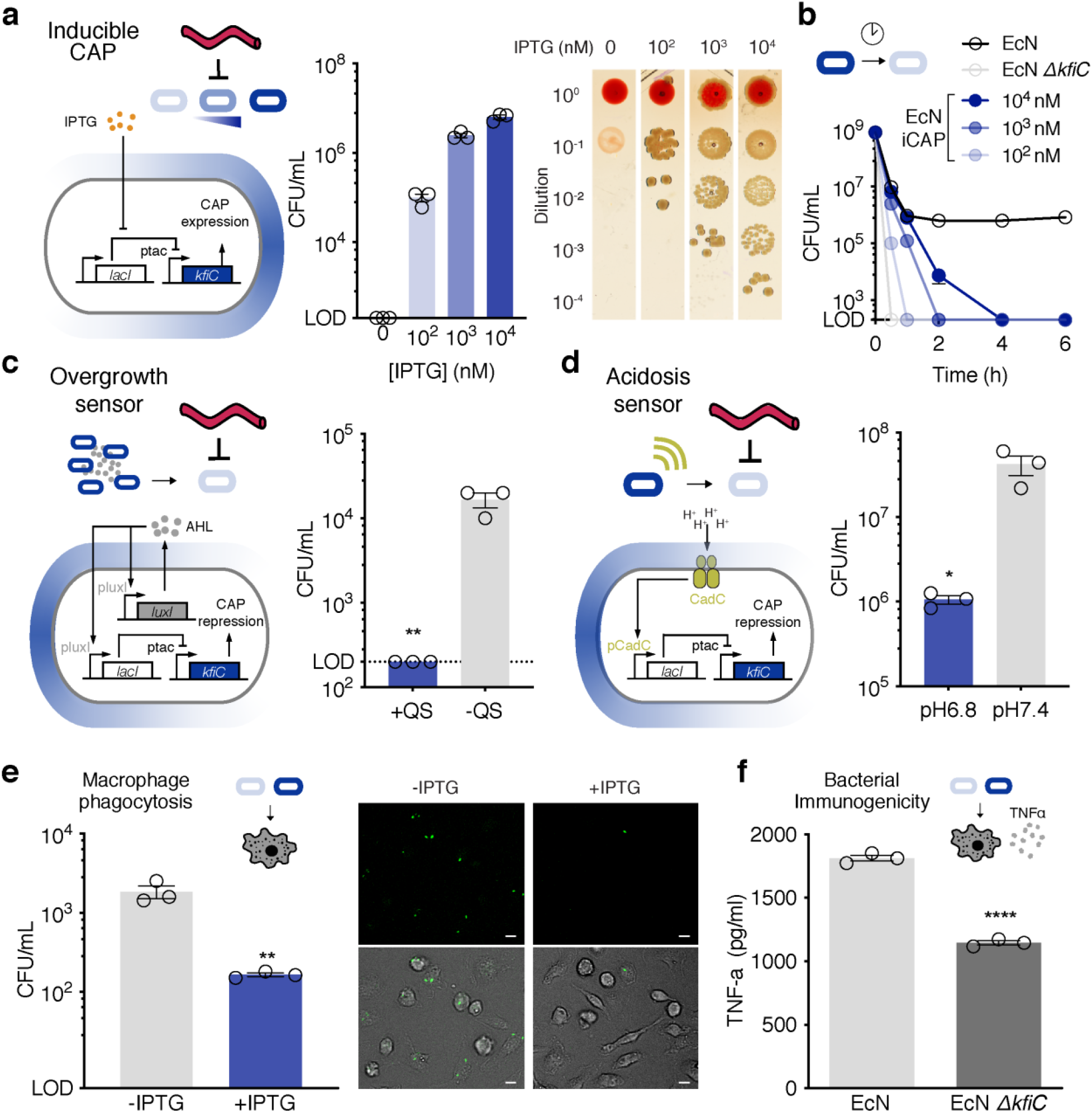
Tunable interaction of the programmable capsular polysaccharides (CAP) system with host immune factors. **a,** Elevating levels of CAP activation with IPTG enabled corresponding increase in bacterial survival in human whole blood (left). Representative images of bacteria spotted on LB-agar plate after 1 hour incubation in human whole blood (right). **b,** Survival kinetics using varying levels of IPTG induction prior to incubation with human whole blood. LOD at 2×10^2^ CFU/mL. **c,** Gene circuit diagram for repression of CAP upon sensing bacterial overgrowth (left). Quorum sensing (QS) *luxI* gene produces diffusible AHL molecules which activates the pluxI promoters. *lacI* gene is placed under the control of pluxI promoter, repressing *kfiC* expression upon reaching quorum. Reduced bacterial survival was achieved after 2 hours inoculation in human whole blood (right; **P = 0.007, unpaired t-test).-QS denotes a control strain with mutated *luxI* gene. LOD at 2×10^2^ CFU/mL. **d,** Gene circuit diagram for repression of CAP upon sensing blood acidosis (left). *lacI* gene is placed under the control of pCadC acid-sensitive promoter, repressing *kfiC* expression upon sensing acidity. Reduced bacterial survival was achieved 4 hours post inoculation in human whole blood at physiologically relevant neutral (pH7.4) and acidic (pH6.8) conditions (right; *P = 0.02, unpaired t-test). **e,** BMDMs were co-cultured with activated or non-activated bacteria for 30 minutes and lysed to enumerate phagocytosed bacterial number. The representative fluorescence microscopy images show decrease in bacteria (GFP, top) in phagocytes (Brightfield, bottom) upon CAP activation (**P = 0.007, unpaired t-test). Scale bar is 10 μm. **f,** CAP contribution to immune recognition. ELISA of TNFα showed a decrease in cytokine production by THP-1 cells incubated with EcN compared to the cells incubated with EcN Δ*kfiC* (****P < 0.0001, unpaired t-test). All error bars represent standard error of mean (SEM) over three independent samples. Limit of detection (LOD) at 2×10^2^ CFU/mL in all panels.

Autonomous systems that repress CAP expression upon sensing of specific conditions would be highly useful to clear bacteria and ensure safety. As a proof of principle, we constructed genetic circuits capable of sensing (1) bacterial overgrowth at colonized sites, and (2) acidosis associated with sepsis^35^ to prevent systemic bacterial growth and inflammation. For both CAP systems, we placed *kfiC* under the control of *tac* promoter on the high (ColE1 origin) copy number plasmid to express CAP despite endogenous *lacI* expression. To design a CAP system responsive to bacterial overgrowth, we incorporated a quorum-sensing module where bacteria express *luxI* gene to produce diffusible small molecule N-Acyl homoserine lactone (AHL). Upon reaching critical population density, the AHL-sensing *pluxI* promoter drives expression of *lacI*, repressing CAP production (Fig. 4c). We cultured the bacteria to stationary phase in LB media to simulate bacterial overgrowth, and observed that this quorum-sensing CAP (qCAP) system resulted in bacterial immunity against ΦK1-5. Control strains harboring a mutated *luxI* gene^36^ were sensitive to the phage, and exogenous addition of AHL molecule rescued bacterial immunity (Supplementary Fig. 8a), confirming CAP repression via quorum-sensing circuit. To test the safety feature of the system, we inoculated the bacteria in human whole blood. Rapid elimination of the qCAP strain was observed after 2 hours while the control strain persisted (Fig. 4c), indicating bacterial overgrowth sensitizes bacteria to immune clearance via qCAP. To sense acidosis, we utilized a previously characterized pH-sensitive promoter pCadC. Membrane tethered endogenous CadC protein is cleaved to activate pCadC promoter in an acidic environment which drives expression of *lacI* to repress CAP production (Fig. 4d). While the bacteria carrying this acidosis-sensing CAP (aCAP) system were sensitive to ΦK1-5 in a physiologically neutral condition (pH 7.4), they were able to grow in the presence of ΦK1-5 in a pH level similar to severe acidosis^37,38^ (pH 6.8) (Supplementary Fig. 8b), indicating CAP repression upon sensing acidity. When the aCAP strain was inoculated in human whole blood, we observed decreased levels of bacteria in acidic condition (Fig. 4d, Supplementary Fig. 8c). These genetic circuits highlight exogenous and autonomous control over bacterial CAP expression, allowing for programmable bacteria sensitivity to host immune detection to enhance safety.

To investigate the effect of the programmable CAP system on bacterial interaction with individual immune factors within whole blood, we assessed how CAP alterations modulated macrophage-mediated phagocytosis and complement-mediated killing. To study phagocytosis, we incubated EcN with murine bone marrow-derived macrophages. iCAP activation prior to co-incubation with macrophages resulted in reduction in uptake of bacteria within macrophages compared to basal control (Supplementary Fig. 9a,b), demonstrating controllable protection from cellular immune recognition. Bacterial colony counting of macrophage lysates and fluorescence microscopy imaging confirmed ~10-fold less phagocytosis with bacteria induced with IPTG compared to uninduced control (Fig. 4e). To assess inflammatory response by the phagocytes, we co-cultured EcN with THP-1 human monocytic cells and measured levels of TNFα, a major cytokine produced in response to microbial detection. Presence of CAP reduced levels of TNFα (Fig. 4f), indicating the ability of CAP to mask microbial recognition from the immune system. To study protection against circulating host antimicrobials such as the complement system, we exposed EcN to human plasma. Presence of CAP improved bacterial survival by at least ~10^5^ fold (Supplementary Fig. 10), demonstrating that CAP protects bacteria from soluble host bactericidal factors. Together, these findings suggest the potential utility of the programmable CAP system to modulates a multitude of host-microbe interactions *in vivo*.

### Transient CAP improves safety and efficacy of engineered probiotic therapy

Intravenous (i.v.) delivery of bacteria allows access to various disease sites in the body; however, systemic delivery of bacteria remains challenging because (1) rapid clearance by the host immune system requires increased dosing, while (2) failure in bacteria clearance can lead to bacteremia and sepsis. Since the programmable CAP system allowed temporal control over bacterial protection and immunogenicity, we sought to improve bacterial delivery by initially protecting bacteria during the delivery stage via CAP production, and subsequently allowing CAP decay to clear them and ensure safety. To study the protective role of CAP *in vivo*, we first characterized probiotic bioavailability and host health in mouse models (Fig. 5a). Upon i.v. administration of EcN Δ*kfiC*, viable bacteria in blood circulation quickly dropped below the LOD (200 CFU/mL). In contrast, EcN remained detectable during the first 4 hours (Supplementary Fig. 11a), demonstrating the protective function of CAP *in vivo*. To examine the host response to encapsulated (*i.e*., wild type) *vs*. unencapsulated (*i.e*., Δ*kfiC*) EcN, we measured levels of serum TNFα and total white blood cell count. We detected lower levels of serum TNFα in the first hour of EcN injection compared to EcN Δ*kfiC* injection (Fig. 5b), similar to the decreased TNFα response observed in our *in vitro* assay. This short-term inflammatory response was resolved within 24 hours for both bacterial strains. In contrast to the rapid resolution of TNFα response, we detected elevated total white blood cell count after 24 hours of injection at higher level with EcN compared to EcN Δ*kfiC* (Supplementary Fig. 11b). Neutrophil expansion accounted for the majority of the immune response to encapsulated EcN (Fig. 5b), suggesting that the persistence of CAP-expressing EcN poses a risk of prolonged bacteremia, which may result in systemic inflammation and toxicity. Thus, while CAP can improve bioavailability, static protection may lead to prolonged bacterial circulation in blood and pose toxicity risks.

**Figure 5:**
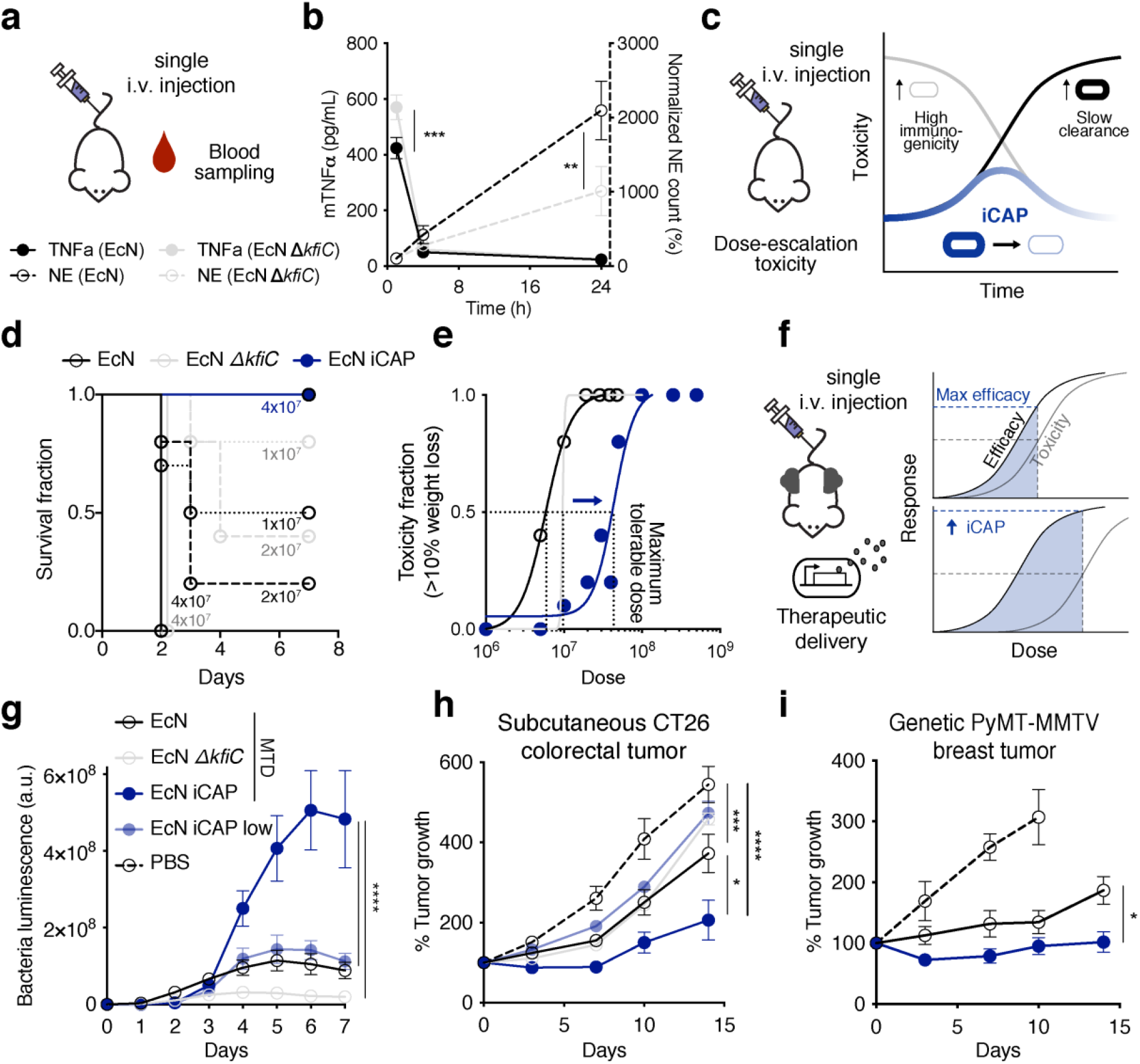
Transient capsular polysaccharides (CAP) activation improves systemic bacterial delivery and efficacy *in vivo*. **a,** Evaluation of host response after bacterial administration. 5×10^6^ bacteria were systemically delivered to BALB/c mice by i.v. injection, and cheek blood was collected at 1, 4, and 24 hours post injection (p.i.). **b,** Change in TNFα levels (solid line) and neutrophil count (dotted line) after bacterial injection. EcN induced lower levels of initial TNFα spikes compared to EcN Δ*kfiC* (***P = 0.0007, two-way ANOVA with Sidak’s multiple comparisons test; n = 5 per group). EcN induced greater levels of neutrophil expansion compared to EcN Δ*kfiC*. Difference in neutrophil levels were observed after 24 hours p.i. (**P = 0.005, two-way ANOVA with Sidak’s multiple comparisons test; n = 5 per group). **c,** Safety evaluation upon i.v. bacteria injection with elevating dosage. BALB/c mice were i.v. administered with EcN iCAP, EcN, or EcN Δ*kfiC* at dosage ranging from 5×10^6^ to 5×10^8^ CFU. EcN iCAP was pre-induced with 10 μM IPTG and allowed for gradual attenuation of CAP over time to minimize toxicity. **d,** Survival curve after bacterial administration. All animals injected with 4×10^7^ CFU of EcN or EcN Δ*kfiC* succumbed within 2 days (>15% body weight reduction). All mice survived after injection with 4×10^7^ CFU of EcN iCAP (n ≥ 5 mice per group). **e,** Dose-toxicity curve. Transient activation of iCAP increased maximum tolerable dose (MTD = 4.4×10^7^ CFU) compared to EcN (5.8×10^6^ CFU) or EcN Δ*kfiC* (9.6×10^6^ CFU). MTD was calculated based on TD50 for exhibiting moderate toxicity (>10% body weight drops p.i.; Nonlinear regression with least squares fit; n ≥ 5 per group). **f,** Therapeutic bacteria administration at MTD to improve antitumor efficacy. Mice bearing tumors were i.v. injected with EcN MTD, EcN Δ*kfiC* MTD, EcN iCAP MTD (pre-induced with 10 μM IPTG), or EcN iCAP low (pre-induced with 10 μM IPTG) expressing antitumor theta-toxin at 5×10^6^, 1×10^7^, 5×10^7^, or 5×10^6^ CFU, respectively. **g,** Bacterial growth trajectories in subcutaneous CT26 tumors after intravenous delivery *in vivo*. Each line represents average of bacterial growth trajectories in tumors quantified by bacterial luminescence over time for each bacterial strain injected. Mice injected with EcN iCAP MTD showed higher bacterial luminescence in tumors compared to mice injected with EcN MTD, EcN Δ*kfiC* MTD, and EcN iCAP low (****P < 0.0001, Two-way ANOVA with Turkey’s multiple comparison test; n = 14, 13, 9, and 13 tumors, respectively, for EcN MTD, EcN Δ*kfiC* MTD, EcN iCAP MTD and EcN iCAP low groups). Luminescence values are normalized to basal luminescence of individual strains. **h,** Therapeutic efficacy measured by relative tumor size over time in a syngeneic CT26 model. EcN iCAP MTD demonstrated highest tumor growth suppression (****P < 0.0001, ***P = 0.0008, **P = 0.003; two-way ANOVA with Tukey’s multiple comparison test; n = 14, 13, 9, 13 and 11 tumors, respectively, for EcN MTD, EcN Δ*kfiC* MTD, EcN iCAP MTD, EcN iCAP low, and PBS groups). **i,** Therapeutic efficacy measured by relative tumor size over time in a genetically engineered spontaneous breast cancer (MMTV-PyMT) mouse model. Tumor growth was measured by calipering three orthotopic regions in mammary glands (upper left, upper right, and bottom). EcN iCAP MTD demonstrated higher tumor growth suppression than EcN MTD (*P = 0.0197; two-way ANOVA; n = 15, 15, and 9 tumors, respectively, for EcN MTD, EcN iCAP MTD, and PBS groups. Mice in PBS groups reached study endpoint 10 days p.i.). All error bars represent SEM.

We hypothesized that transient activation of the programmable CAP system can improve bacterial delivery profiles by modulating maximum injectable dose, host toxicity, and biodistribution. Inducing CAP expression prior to injection would improve bioavailability and mask cytokine induction, and loss of CAP in the absence of the inducer *in vivo* would effectively clear bacteria and minimize long-term immune responses. To test this strategy, we first i.v. administered escalating doses of EcN iCAP and assessed host health and determined maximum tolerable dose^39,40^ (MTD) (Fig. 5c). At lower doses, EcN iCAP caused a smaller decrease in body weight compared to EcN and EcN Δ*kfiC* with static cellular surface (*i.e*., with or without CAP, respectively) (Supplementary Fig. 12a,b). Importantly, EcN iCAP dramatically reduced toxicity compared to EcN and EcN Δ*kfiC* at higher doses: EcN and EcN Δ*kfiC* caused severe end-point toxicity (death or >15% loss of weight) to mice treated with doses above 1×10^7^ CFU within 2 days, while no mice showed severe toxicity following injection of pre-induced EcN iCAP at the same doses (Fig. 5d). Based on these data, we computed a dose-toxicity curve and demonstrated that transiently induced EcN iCAP results in ~10-fold higher MTD compared to EcN and EcN Δ*kfiC* (Fig. 5e). To further study safety, we simulated a severe toxicity scenario by inducing sepsis by intraperitoneal injection of bacteria^41^. At both high and low doses (10^7^ and 10^6^ CFU, respectively), we consistently observed improved safety for EcN iCAP compared to EcN and EcN Δ*kfiC* (Supplementary Fig. 13a,b). Finally, to study bacteria biodistribution, we administered all groups of EcN at a matched dose of 5×10^6^ CFU via i.v. injection. Approximately 10-fold less EcN and EcN iCAP were found in peripheral organs (liver and spleen) compared to EcN Δ*kfiC* (Supplementary Fig. 14), indicating that initial induction of EcN iCAP was sufficient to provide protection from the mononuclear phagocyte system. These data support that transient activation of iCAP improves probiotic delivery and safety.

Since systemic bacterial delivery has been extensively used for cancer therapy, we next tested whether the programmable CAP system can improve antitumor efficacy by permitting higher doses (Fig. 5f). To engineer bacteria to deliver antitumor payloads, we cloned a gene encoding pore-forming toxin, theta toxin (TT), previously shown to be effective as a bacterial cancer therapy^42^, in a high copy number plasmid (ColE1) with a stabilization mechanism for *in vivo* applications (Axe/Txe system^43^). In a syngeneic CT26 colorectal cancer model, we intravenously administered engineered EcN at the corresponding MTD of each strain (EcN, EcN Δ*kfiC*, and EcN iCAP at 5×10^6^, 1×10^7^, and 5×10^7^ CFU, respectively), along with a low dose of EcN iCAP at 5×10^6^ CFU to match the MTD of EcN. Over the following days, bacterial accumulation in tumors was observed by luminescence. Here, EcN iCAP MTD showed significantly higher signals in tumors compared to all other groups (Fig. 5g). After bacterial administration, we observed that mice treated with EcN MTD, EcN Δ*kfiC* MTD, and low dose EcN iCAP exhibited modest tumor growth suppression compared to untreated group over 14 days. By contrast, single administration of EcN iCAP MTD resulted in significant tumor growth suppression by ~400% compared to the untreated group (Fig. 5h). While increased MTD enabled by transient activation of the iCAP system improved therapeutic efficacy, body weight of animals between MTD groups remained similar (Supplementary Fig. 15). We next compared efficacy of TT-producing EcN and EcN iCAP at MTD in a genetically engineered spontaneous breast cancer model (MMTV-PyMT). EcN iCAP MTD resulted in improved tumor growth suppression by ~100% compared to EcN MTD over 14 days (Fig. 5i). Consistently, we observed higher bacterial signal in tumor from EcN iCAP compared to EcN while body weight remains similar between the two treatment groups (Supplementary Fig. 16). To further explore the role of CAP on bacterial delivery to tumors *in vivo*, we built a mathematical bacterial pharmacokinetics model. Our simulations suggested transient protection of bacteria using the iCAP system could improve tumor specificity by minimizing persistence in peripheral organs (*i.e*., blood and liver), supporting our experimental observations (Supplementary Fig. 17a). As a result, this approach allows for elevated bacterial doses, which leads to increased tumor accumulation upon injection. Taken together, the iCAP system enables increased tolerable bacterial doses and improved therapeutic efficacy.

### In situ CAP activation translocates EcN to distal tumors

Intratumoral (i.t.) bacteria injection has been used as a route of delivery in clinical settings due to higher therapeutic efficacy, dose titration capability, and improved safety profiles compared to systemic injection^44–48^. One unique capability of i.t. delivery is the translocation of bacteria from injected tumors to distal tumors^48^, potentiating a novel route of safe bacterial delivery to inaccessible tumors. However, continuous translocation coupled with long-term survival of bacteria can pose a significant safety concern; thus transient *in situ* activation could allow for more optimal utilization of this phenomena. To model this hypothesis, we simulated i.t. delivery and showed that *in situ* induction of EcN iCAP within the tumor increases bacterial bioavailability in circulation and facilitates bacterial translocation to distal tumors (Fig. 6a, Supplementary Fig. 17b). We then tested this strategy via i.t. injection of uninduced EcN iCAP (*i.e*., without CAP) into a single tumor of mice harboring dual hind-flank CT26 tumors (Fig. 6b). To activate the iCAP system *in situ*, mice were fed with water containing IPTG. After 3 days, we observed a marked increase in bacterial translocation to distal tumors compared to uninduced bacteria (Fig. 6c, Supplementary Fig. 18a,b). Biodistribution data showed tumor-specific translocation (Fig. 6d, Supplementary Fig. 18c), and mice exhibited minimal reductions in body weight (Supplementary Fig. 18d). Tracing colonization kinetics using bioluminescent EcN confirmed the appearance of bacteria in distal tumor 1 day following IPTG administration (Supplementary Fig. 19a). We next explored whether this bacterial trafficking approach can be generalized in multiple clinically-relevant animal models. We tested orthotopic breast cancer (mammary fat-pad 4T1) and MMTV-PyMT mouse models. Consistently, we observed increased bacterial translocation to distal tumors via *in situ* activation of iCAP in both tumor models (Fig. 6c, Supplementary Fig. 1b,c, Supplementary Fig. 20a-c, Supplementary Fig. 21a-c). Notably, i.t. injection of EcN iCAP into a single tumor in the MMTV-PyMT model resulted in microbial translocation to multiple distal tumors throughout the body following IPTG induction. These results demonstrate the robustness of iCAP-mediated translocation across a range of locations and tumor types.

**Figure 6:**
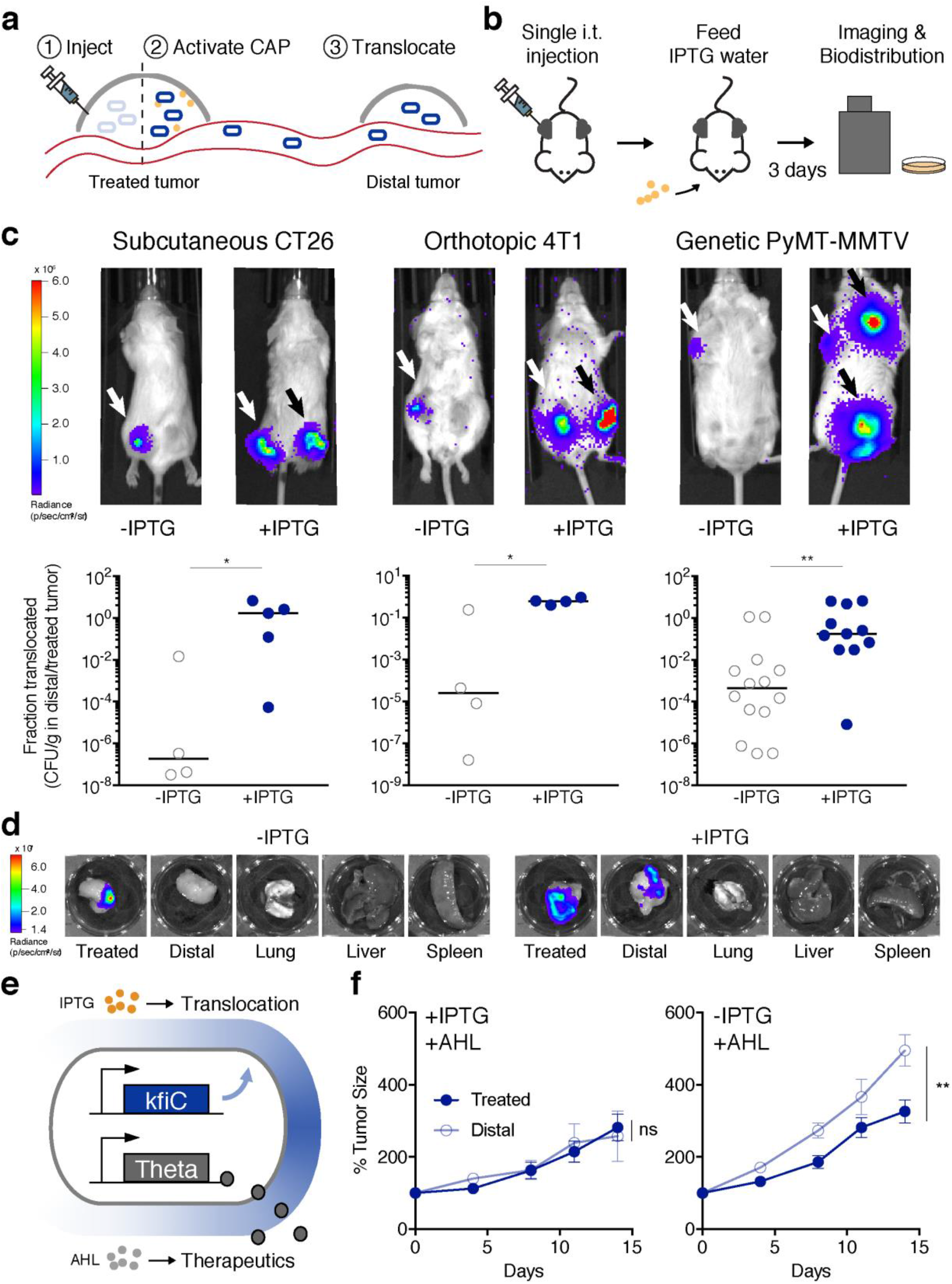
*In situ* activation of the programmable capsular polysaccharides (CAP) enables bacterial translocation and drug delivery to distal tumors. **a,** Schematics of iCAP-mediated bacterial translocation. EcN are injected into one tumor (treated tumor). iCAP activation enables bacteria translocation to distal tumors. **b,** Mice harboring multiple tumors are injected with EcN iCAP into a single tumor (treated tumor). Subsequently, mice are fed 10 mM IPTG water to activate iCAP *in situ*. Mice are imaged daily for bacterial bioluminescence to track tumor colonization in tumors. To quantify bacterial biodistribution, organs are harvested and bacterial colonies are counted after 3 days. **c,** Inducible translocation of EcN iCAP to distal tumors in CT26 syngeneic (left), 4T1 orthotopic (middle), and MMTV-PyMT genetically engineered (right) mouse tumor models. Representative IVIS images showing bacterial translocation *in vivo*. White arrows indicate location of bacterial injection. Black allows indicate location of bacterial translocation. Translocation is quantified by fraction of bacteria found in distal tumor compared to treated tumor. Bacteria number is measured by performing biodistribution for CFU/g enumeration. iCAP activation showed marked increase in bacterial translocation (*P = 0.032, *P = 0.029, **P = 0.003, Mann-Whitney test). **d,** Representative images of *ex vivo* organ images taken with IVIS showing bacterial tumor translocation in 4T1 orthotopic mouse model. **e,** Schematics of engineered EcN capable of programmable translocation and therapeutic expression. Therapeutic production is externally controlled by an inducer AHL. The engineered EcN was injected into a single tumor and IPTG-induced to translocate to distal tumors, and AHL-induced to deliver therapeutics. **f,** Therapeutic efficacy in treated and distal CT26 tumors measured by relative tumor growth over time. Bacteria were injected into a single treated tumor. The translocation was controlled by IPTG water. 3 days p.i., AHL was administered to induce therapeutic expression. Distal tumor growth was suppressed only when therapeutic bacteria were able to translocate (n.s. P = 0.83, **P = 0.004, two-way ANOVA with Bonferroni posttest, n = 6, 5 for both treated and distal tumors, respectively). All error bars represent SEM.

We next delivered therapeutics to tumors using engineered EcN expressing the antitumor TT payload. TT was cloned under the *luxI* promoter that is responsive to an inducer molecule AHL orthogonal to IPTG (Fig. 6e). Following i.t. injection of therapeutic EcN into a single tumor in the CT26 dual flank mouse model, translocation to uninjected tumors was controlled by feeding mice with or without IPTG water. Another group of mice were given i.t. injection of EcN iCAP without TT as non-therapeutic control. iCAP-mediated bacterial translocation was confirmed by bacterial bioluminescence (Supplementary Fig. 22a-c). Subsequently, AHL was administered subcutaneously to induce TT expression, and tumor growth was monitored. While PBS treatment allowed tumor growth of ~400%, a reduction in the growth of tumors were observed when they were directly injected with therapeutic EcN regardless of bacterial translocation. By contrast, therapeutic efficacy in distal (uninjected) tumors was only observed when the mice were fed with IPTG water followed by TT induction via subcutaneous injection of AHL (Fig. 6f, Supplementary Fig. 23), demonstrating successful therapeutic delivery to distal tumors using the iCAP-mediated bacterial translocation approach. Body weight quickly recovered to baseline within a few days post administration for all conditions (Supplementary Fig. 24). Together, we provide a demonstration of controllably translocating therapeutic bacteria and utilizing this strategy to treat distal tumors *in vivo*.

### Conclusion

We have demonstrated a synthetic biology approach for dynamic and tunable modulation of the bacterial surface in the context of *in vivo* therapeutic delivery. Several biosensing circuits were designed to allow for exogenous and autonomous regulation of CAP expression, which were shown to enhance both safety and efficacy in multiple therapeutically-relevant scenarios. Taking advantage of the natural evolution of the CAP system to interface with a multitude of environments, we showed engineered bacterial interactions with host immunity, bacteriophage, antimicrobials and acidity. Though we have explored the modulation of CAP density via *kfiC* in this study, additional genes identified through our sRNA screen may also be used to achieve varied sensitivities or to independently alter bacterial sensitivity to various environmental factors. For example, *kfiB* and *kfiC* genes have been reported to differentially modulate bacterial interactions with epithelial cells^29^. We have also shown that this approach can be applied to other strains of *E. coli*. Since there exists over 80 distinct *E. coli* CAP systems^24,31^ and many more in other species, we envision CAP engineering to possess vast opportunities to controllably modulate microbial surface properties for therapeutic delivery.

Despite recent preclinical progress with bacterial therapies, dose-limiting toxicity has been a long-standing challenge, slowing efforts for clinical translation. Early works by William Coley in the 19^th^ century had observed tumor regression upon injections of a live bacterial cocktail^49^, but this approach was largely unsafe due to potential risks of infections and inflammatory side effects. More recently, several clinical trials using genetically attenuated bacterial strains observed dose-limiting toxicities. For example, systemic administration of attenuated *Salmonella typhimurium* strain (VNP20009) at or higher than tolerable dose led to tumor colonization in < 20% of patients, and resulted in no objective regression in a phase I clinical trial^11^. While the recent focus of synthetic biology has been the engineering of various therapeutic payloads to increase efficacy, strategies to improve bacterial delivery has been limited. We utilized on-demand CAP system to controllably protect probiotic EcN from immune clearance, and demonstrated an ~10-fold increase in the systemically injectable tolerated dose *in vivo*. As a result, transient expression of CAP was able to safely enhance bacterial delivery and suppress rapidly growing syngeneic colorectal tumors with a single i.v. administration. Given that humans are 250-fold more sensitive to endotoxins compared to mice^50^, we expect our results to have important implications in the clinical translation of bacterial therapies. Since the CAP system is orthogonal to other bacterial surface structures such as LPS, phospholipids, and flagellum^51^, synergistic combinations of these systems could further improve the programmability and immunogenicity of bacterial therapy, and warrant future investigation.

We further devised the programmable CAP system to facilitate translocation of bacteria to distal tumors, demonstrating a novel delivery strategy with advantageous safety profiles. Comparing across different tumor types, we noted leakier translocation of bacteria in 4T1 and PyMT-MMTV models, suggesting tumor type, location, or vascularization may play a role in bacterial escape from the tumor microenvironment. Regardless, *in situ* activation of the programmable CAP system opens the possibility of utilizing programmable translocation of therapeutic bacteria to inaccessible disease sites including metastatic tumors. Transient *in situ* activation could allow for colonization of newly formed tumors with reduced accessibility. As humarns are more sensitive to bacteria compared to mice, we expect that the programmable CAP system will be far more effective in human than mice at controlling bioavailability of bacteria in circulation. In addition, the ability to transiently increase circulating bacteria from colonized tumor may allow safe sampling of bacteria without the need for an invasive tumor biopsy while preventing excessive leakage of bacteria that could cause bacteremia and systemic infection. More broadly, we envision that *in situ* control over bacterial CAP may be utilized to provide further safeguards during bacterial therapy such as sensitization of bacteria with antibiotics^52,53^ and phage therapy^54^.

In addition to clear applications within bacteria cancer therapy, the utility of the programmable CAP system can be extended to other clinical settings. For example, we showed that CAP provides protection from acids which could potentially protect orally-delivered probiotics during transit through the gastric environment and facilitate intestinal colonization^55^. Beyond delivery, altering surface immunogenicity may also be utilized therapeutically to modulate the host immune landscape in the tumors and gut^56,57^. The platform described here could also be used as a model system to study the role of CAP and other surface structures on pathogen colonization in the host environment^58,59^. Aside from model systems and clinical applications, the programmable CAP system may provide a general platform for a programmable interface with various environments. As microbial deployment in various applications continue to advance, robust control over microbial interaction with complex surroundings will ensure safe and effective implementation of engineered microbes.

## Acknowledgments

We thank Dr. Ian Molineux for providing ΦK1-5 phage, and K1 and K5 *E. coli* strains. We thank Dr. Kunihiro Uryu at EMSCOPIC for his technical support.

## Funding

This work was supported by the NIH Pathway to Independence Award (R00CA197649–02), DoD LC160314 (T.D.), DoD BC160541 (T.D.), NIH R01GM069811 (T.D.), NIH F99CA253756 (T.H.), and Honjo International Foundation Scholarship (T.H.).

## Author contributions

T.H., J.H., K.L. and T.D. conceived and designed the study. T.H., J.H., Y.C., J.I., J.Z., F.L., S.C. and C.N. performed *in vitro* characterization. T.H., J.H., Y.C., C.C., K.G., N.H. and K.P. performed *in vivo* experiments for this study. T.H., J.H., N.H. and T.D. developed computational modeling for this study. T.H., J.H., N.A., K.L. and T.D. wrote the manuscript.

## Competing interests

T.H., J.H., K.L. and T.D. have filed a provisional patent application with the US Patent and Trademark Office related to this work.

## Data and materials availability

All data is available in the main text or the supplementary materials.

## Materials and Methods

### Bacterial strains and culturing

The host strain used in this study was *Escherichia coli* Nissle 1917 (EcN) that naturally expresses K5 capsular polysaccharide (CAP) containing a genomically integrated erythromycin-resistance *luxCDABE* cassette for bacterial bioluminescence tracking *in vivo*. For all strains used in this study, please refer to Table S1. All bacteria were grown with appropriate antibiotics selection (100 μg/mL ampicillin, 50 μg/mL kanamycin, 25 μg/mL chloramphenicol, 50 μg/mL erythromycin) in LB media (Sigma-Aldrich) at 225 RPM or on LB-agar plates containing 1.5% agar at 37°C.

### Construction of plasmids and gene circuits

To construct a knockdown library, plasmids with sRNA targeting each gene of the CAP biosynthetic pathway were prepared using Gibson Assembly. The sRNA sequences were designed to be complementary and bind to the 24-neucleotide sequence of the target gene coding sequence spanning the ribosome binding site and the start codon^1,2^. A plasmid template was prepared by PCR-amplifying backbone (pTH05) using primers (pTH05_for and pTH05_rev), and the single-stranded DNA for sRNA against genes in CAP biosynthesis were inserted (*kfiA*, *kfiB*, *kfiD*, *kpsC*, *kpsS*, *kpsF*, *kpsU*, *kpsE*, *kpsD*, *kpsT*, *kpsM*), and transformed into Mach1 competent cells (Invitrogen). CAP gene circuits and the therapeutic plasmids were constructed in a similar manner. Genes of interest were obtained by synthesizing oligos or gBlock from IDT, or PCR-amplification (*kfiC* gene was obtained via colony PCR from EcN). Subsequently, plasmids were constructed using Gibson Assembly or using standard restriction digest and ligation cloning, and transformed into Mach1 competent cells (Invitrogen).

### Construction of knockout strains

EcN was transformed to carry Lambda Red helper plasmid (pKD46)^3^. Transformants were grown in 50 mL LB at 30°C with chloramphenicol to an OD600 of 0.4 and made electrocompetent by washing three times with ice cold MilliQ water and concentrating 150-fold in 15% glycerol. Chloramphenicol-resistance cassette was prepared by PCR with primers flanked by sequence within each target gene followed by gel purification and resuspension in MilliQ water. Electroporation was performed using 50 μL of competent cells and 10-100 ng of DNA. Shocked cells were added to 1mL SOC, incubated at 30°C for 1 hour with 20 μL arabinose, and incubated at 37°C for 1 hour. Cells were then plated on LB plates with chloramphenicol and incubated in 37°C overnight. Colonies were picked the next day to obtain knockout strains including Δ*kfiC* strain (EcN Δ*kfiC*).

### Characterization of CAP strains sensitivity to phages, antibiotics, and acids

To perform plaque forming assay, bacteria were plated onto LB agar plates to make a lawn and allowed to dry under fire. 10 μL of serial diluted ΦK1-5 phage (Molineux, University of Texas, Austin) was spotted onto the plates and allowed to dry. Plates were incubated at 37°C overnight and inspected the next day for plaque forming unit (PFU) counting. Similar phage plaque forming assay were performed for K1 and K5 type *E. coli* strains.

To assess bacterial growth in liquid culture, overnight cultures of EcN, EcN Δ*kfiC*, or EcN iCAP strains were calibrated into OD600 of 1.0, and 100 μL of each was transferred into 96-well plate (Corning). 1 μL of 10^8^ PFU ΦK1-5 phage, or antibiotics of indicated concentrations were added to each well. The samples were incubated at 37°C with shaking in Tecan plate reader, and the OD600 was measured every 20 min. For bacterial growth in low pH condition, LB media adjusted to pH2.5 using HCl, bacteria were incubated at 37°C for 1 hour, followed by serial dilution and plating on a LB agar plate for CFU enumeration.

### Characterization of CAP using SDS-PAGE

CAP was purified via the chloroform-phenol extraction as previously described^4,5^. Briefly, 3 mL of overnight bacteria cultures were harvested the next day and further sub-cultured in 50 mL LB broth in the presence or absence of 0.1 M IPTG for indicated lengths of time. Bacteria concentrations were adjusted to the same level across samples via OD600 before centrifugation. Pellets were collected and resuspended in 150 μL of water. An equal amount of hot phenol (65°C) was added, and the mixtures were vortexed vigorously. The mixtures were then incubated at 65°C for 20 minutes, followed by chloroform extraction (400 μL) and centrifugation. The CAP were detected by alcian blue staining as previously reported^5–7^. Briefly, following SDS-PAGE electrophoresis (4-20% gradient), the gel was fixed in fixing solution (25% ethanol, 10% acetic acid in water) for 15 minutes while shaking at room temperature. The gel was then incubated in alcian blue solution (0.125% alcian blue in 25% ethanol, 10% acetic acid in water) at room temperature for 2 hours while shaking before de-stained overnight in fixing solution. CAP was visualized as alcian blue stained bands on the resulting gel.

### Visualization of CAP using TEM

Bacteria were grown overnight in LB media with appropriate antibiotics before being processed for imaging. For EcN iCAP, a 1:100 dilution in LB with antibiotics was made the following day and grown in 37°C shaker until OD600 = 0.1–0.4 (mid-log phase), and varying concentrations of IPTG were added for further incubation for 6 hours before being processed. The cultures were spun down at 300 relative centrifugal force (rcf) for 10 min and embedded in 2% agarose. Each agarose gel fragment was cut into a cube with 2-mm edge and placed in a 1.5-mL centrifuge tube. The samples embedded in agarose were fixed and stained via protocols previously reported^8^. Briefly, the samples were fixed with 2% paraformaldehyde and 2.5% glutaraldehyde in osmotically adjusted buffer (0.1 M sodium cacodylate, 0.9 M sucrose, 10 mM CaCl_2_, 10 mM MgCl_2_) with 0.075% ruthenium red and 75 mM lysine acetate for 20 min on ice. The samples were washed with osmotically adjusted buffer containing 0.075% ruthenium red twice and further fixed with 1% osmium tetroxide in osmotically adjusted buffer containing 0.075% ruthenium red for an hour on ice. The samples were washed three times in water with 5 min incubation between each wash and dehydrated in increasing concentrations of ethanol (50%, 70%, and 100%) on ice for 15 min per step. The samples were washed one more time in 100% ethanol and embedded in increasing concentrations of Spurr’s resin (33% and 66%) diluted in ethanol for 30 min per step and overnight in 100% Spurr’s resin. The samples were moved to fresh Spurr’s resin the next day and polymerized at 65°C overnight before sectioned using SORVALL MT-2B Ultramicrotome to ~70 nm. The sample sections were placed on TEM grids (Ted Pella; 01800F) and stained using UranyLess (EMS). The sample grids were imaged using FEI Talos 200 TEM.

### TEM image processing and data analysis of polysaccharide layer

The image processing of TEM images was performed using ImageJ, and the data analysis was done using MATLAB. Due to low signal-to-noise ratio of the TEM images resulting from thinly sectioned bacteria samples stained using ruthenium red, Gaussian blur was used to reduce the noise and help determine boundary of polysaccharide layer. Polysaccharide layer was selected and transformed into binary image using threshold function. Some portion of boundary of polysaccharide layer was manually outlined when the thresholding function could not determine where the boundary is. The resulting binary image of polysaccharide layer was used to identify the centroid and measure distribution of polysaccharide thickness in respect to the centroid. For each sample, five representative images were used to measure the polysaccharide thickness and the measurements were aggregated to form histograms. The resulting histograms were fitted with Gaussian curves to extract mean and standard deviation of polysaccharide layer thickness.

### In vitro whole blood bactericidal assays

EcN, EcN Δ*kfiC*, or EcN iCAP bacterial cultures were grown overnight in LB broth with appropriate antibiotics and IPTG concentrations. The cultures were spun down at 3000 rcf for 5 min and resuspended in 1 mL sterile PBS. They were further normalized to an OD600 of 1 with sterile PBS. 150 μL of blood from the single donor human whole blood or murine (BALB/c) whole blood (Innovative Research) were aliquoted into 3 wells/strain in a 96-well plate. 1.5 μL of bacteria were added to each well and incubated at 37°C. At various time points, the plate was taken out, and a serial dilution of each sample was prepared in PBS. The dilutions were plated on LB agar plates with erythromycin. The agar plates were incubated at 37°C overnight and inspected the next day for CFU counting.

### Phagocytosis assays

The phagocytosis assays were performed via protocols as previously reported^9,10^. Briefly, bone marrow derived macrophages (BMDM) were thawed on a 15 cm non-TC treated petri dish and cultured in RPMI with 10% FBS and MCSF for 4 days before experiment. On the 4^th^ day, BMDMs were collected, counted and diluted to 2 × 10^5^ cells/mL in RPMI with 10% FBS (without antibiotics). Afterwards, 1 mL of the new mixture was plated per well (2 × 10^5^ cells) in a 24-well TC-treated plate and cultured overnight. Media in the 24-well BMDM plate was removed the next day, and 1 mL of EcN iCAP constitutively expressing GFP with or without IPTG induction were resuspended in RPMI with 10% FBS without antibiotics was added into each well at MOI of 100. The co-culture was incubated for 30 min at 37°C followed by rigorous washing with PBS at least 3 times. 1 mL of RPMI with 10% FBS and gentamicin (30 μg/mL) was added to each well, followed by live imaging under confocal microscopy. 0.1 M IPTG was added to EcN iCAP with IPTG induction the entire time. Then, the BMDMs were lysed with 0.5% TritonX in PBS and lysates were collected and plated on LB agar with erythromycin followed by overnight incubation at 37°C. Colonies were counted the next day. ImageJ was used to count the number of macrophages, engulfed bacterial cells, and macrophages containing engulfed bacterial cells from the confocal images. The phagocytic index was calculated according to the following formula: phagocytic index = (total number of engulfed bacterial cells/total number of counted macrophages) × (number of macrophages containing engulfed bacterial cells/total number of counted macrophages) × 100.

### Determination of TNF-alpha response

THP-1 cells (ATCC) were maintained in RPMI-1640 supplemented with 10% FBS, 2 mM L-glutamine, 100 μg/mL streptomycin, 100 μg/mL penicillin, and 0.1% mercaptoethanol at 37°C and 5% CO_2_. Cells were passaged every 72 hours. For cell quantification and viability analysis, cells were stained using trypan blue stain. For *in vitro* TNF-alpha assay, THP-1 was resuspended at a concentration of 1 × 10^6^ cells/mL in RPMI-1640 supplemented with 10% FBS and 0.1% gentamycin. 300 μL of cell suspension was transferred into each well of a 24-well plate. 3 μL of each bacterial strain at each concentration were added to cell culture wells. Subsequently, the culture medium was harvested and centrifuged at 200 rcf for 5 min to isolate THP-1 without causing cell death. Supernatant was then centrifuged at 3000 rcf for 5 min to remove bacteria. The resulting supernatant was analyzed for TNF-alpha response. TNF-alpha was measured using an R&D Systems Quantikine ELISA Kit in a plate reader.

### Animal models

All animal experiments were approved by the Institutional Animal Care and Use Committee (Columbia University, protocols ACAAAN8002 and AC-AAAZ4470). For tumor-bearing animals, euthanasia was required when the tumor burden reaches 2 cm in diameter or after recommendation by the veterinary staff. Mice were blindly randomized into various groups. Animal experiments were performed on 8–12 weeks-old female BALB/c mice (Taconic Biosciences). Tumor models were established with bilateral subcutaneous hind flank injection of mouse colorectal carcinoma CT26 cells (ATCC) or mammary fat pad injection of 4T1-luciferase mammary carcinoma cells (Kang, Princeton University). The concentration for implantation of the tumor cells was 5×10^7^ cells per ml in RPMI (no phenol red). Cells were injected at a volume of 100 μl per flank, with each implant consisting of 5 × 10^6^ cells. Female transgenic MMTV-PyMT mice (Jackson Laboratory) which develops mammary tumors were also used. Tumors were grown to an average of approximately 200–400 mm^3^ before experiments. Tumor volume was quantified using calipers to measure the length, width, and height of each tumor (V = L × W × H). Because z dimension of PyMT tumor is highly variable, total volume was calculated as length × width^2^ × 0.5. Volumes were normalized to pre-injection values to calculate relative or % tumor growth on a per mouse basis.

### Bacterial administration for in vivo experiments

Overnight cultures of EcN, EcN Δ*kfiC*, and EcN iCAP were grown in LB medium with the appropriate antibiotics and inducers. A 1:100 dilution in LB with appropriate antibiotics and inducers was made the following day and grown in 37°C shaker until OD600 = 0.1–0.4 (mid-log phase). Cultures were centrifuged at 3000 rcf for 10 min and washed three times with cold sterile PBS. The bacteria were then normalized to a desired OD600. Unless otherwise noted, intravenous injections were given through the tail-vein at the dose of 5 × 10^6^ cells/mL (OD600 of 0.5) in PBS with a total volume of 100 μL per mouse. Intratumoral injections of bacteria were performed at a concentration of 5 × 10^6^ cells/mL with a total volume of 40 μL per tumor. Intraperitoneal injections were injected at varying concentrations in PBS with a total volume of 100 μL per mouse. For induction of theta toxin production, AHL subcutaneous injection was given to mice daily at 10 μM concentration with a total volume of 500 μL per mouse. For *in situ* activation of iCAP, water containing 10 mM IPTG was given to mice a day after bacterial administration.

### Biodistribution and in vivo animal imaging

All bacterial strains used in this study had integrated *luxCDABE cassette* that could be visualized by IVIS spectrum imaging system (Perkin Elmer) and were quantified by Living Image software. Images and body weight of mouse were obtained every day starting the day of bacterial administration until the study endpoint. At the study endpoint, mice were euthanatized by carbon dioxide, and the tumors and organs (spleen, liver, and lungs) were extracted and imaged. They were later weighed and homogenized using a gentleMACS tissue dissociator (C Tubes, Miltenyi Biotec). Homogenates were serially diluted with sterile PBS and plated on LB agar plates with erythromycin and incubated overnight at 37°C. Colonies were counted the next day.

### Statistical analysis

Statistical tests were performed either in GraphPad Prism 7.0 (Student’s t-test and ANOVA) or Microsoft Excel. The details of the statistical tests are indicated in the respective figure legends. When data were approximately normally distributed, values were compared using either a Student’s t-test, one-way ANOVA for single variable, or a two-way ANOVA for two variables. Mice were randomized into different groups before experiments.

