## Supplementary Materials for "A programmable probiotic encapsulation system enhances therapeutic delivery *in vivo*"

### Supplementary Figures and Tables

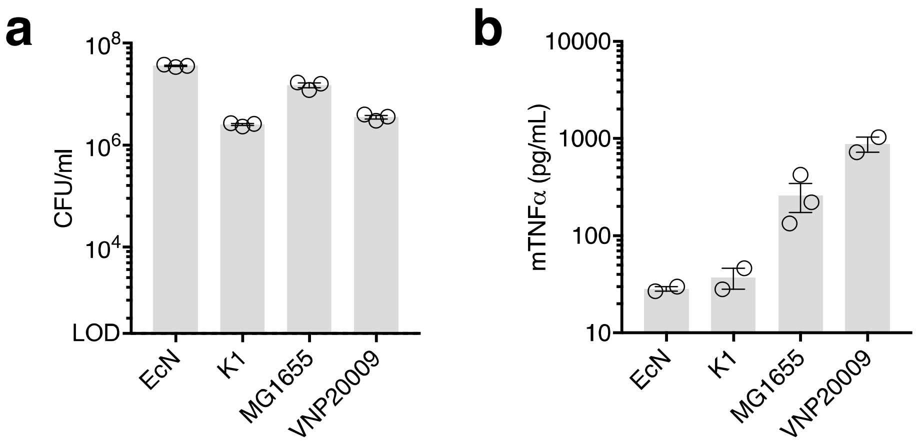


**Supplementary Figure 1: Survival and immunogenicity of *E. coli* and *S. typhimurium* strains. a,** Bacterial survival in human whole blood. $10^8$ CFU/mL bacteria was incubated in the blood for 2 hours and plated on LB agar for CFU enumeration. **b,** Immune recognition of the bacteria. TNFα production by THP-1 cells co-cultured with bacteria for 0.5 hour, measured by ELISA. All error bars represent standard error of mean (SEM).

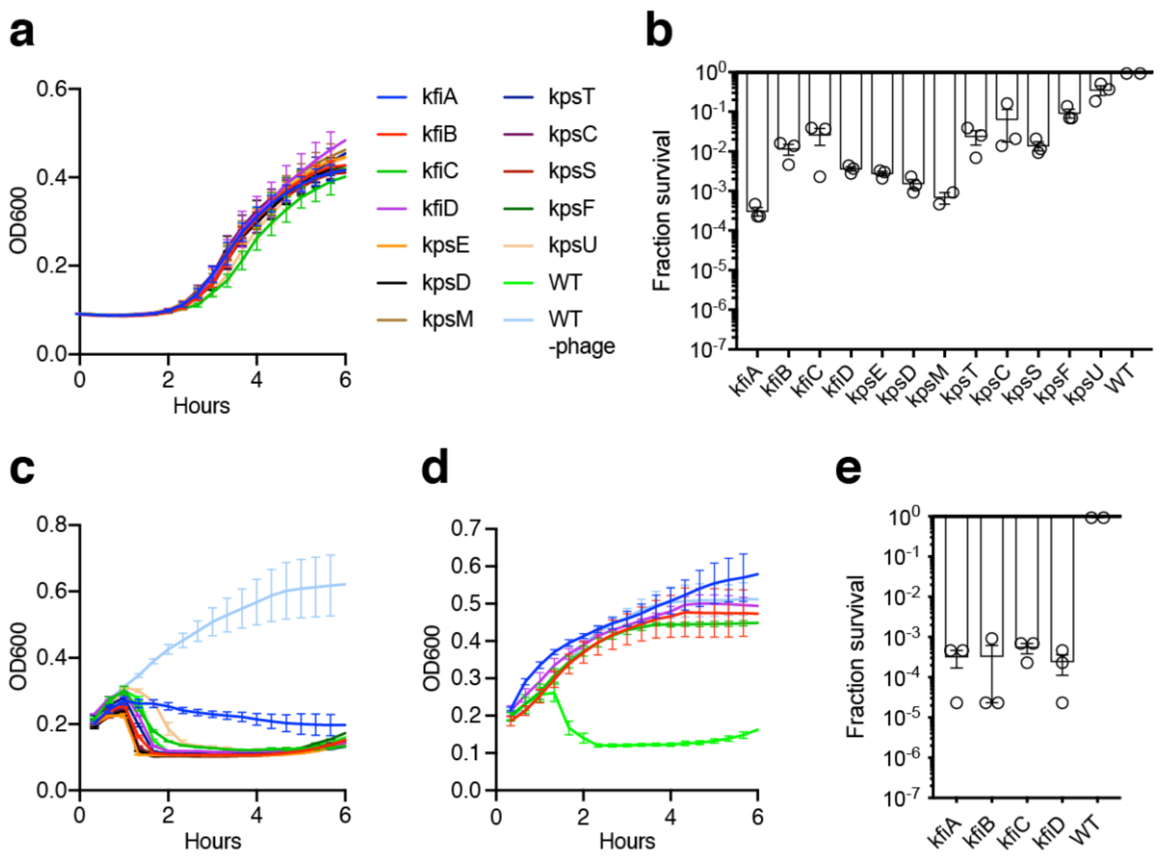


**Supplementary Figure 2: Characterization of sRNA knockdown (KD) and knockout (KO) strains. a,** Growth kinetics of KD strains of *E. coli* Nissle 1917 (EcN) in LB media. $OD_{600}$ was measured over time in a plate reader. **b,** KD strain survival in human blood. Bacteria were inoculated in human whole blood for 0.5 hour, and plated on LB agar for CFU enumeration. Survival fraction is fraction of CFU of KD strain over CFU of wild-type (WT) strain. **c,** Growth of KD strains in LB media containing ΦK1-5. WT strain without ΦK1-5 was included as a baseline bacterial growth. **d,** Growth of KO strains in LB media containing ΦK1-5. WT strain without ΦK1-5 was included as a baseline bacterial growth. (n = 3 per group. All error bars represent SEM.) **e,** KO strain survival in human blood. Bacteria were inoculated in human whole blood for 0.5 hour and plated on LB agar for CFU enumeration. Survival fraction is fraction CFU of KD strain over CFU of WT strain.

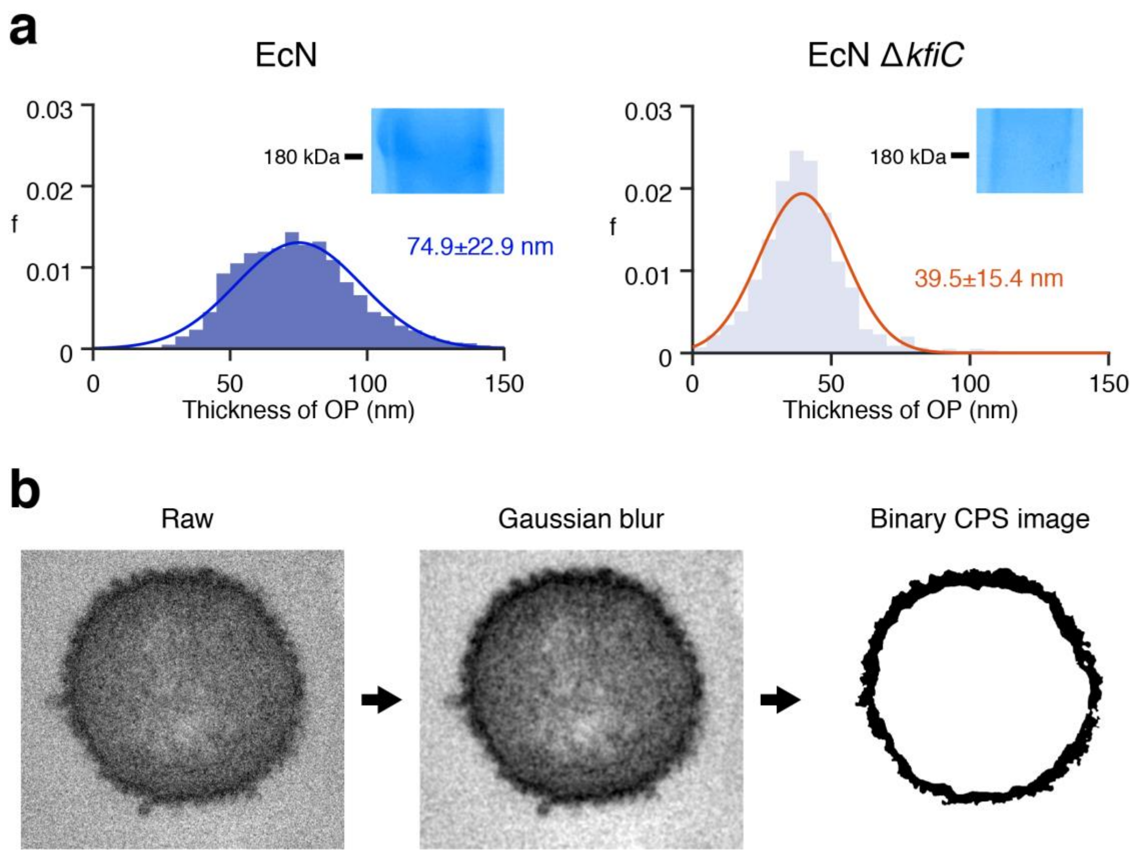


**Supplementary Figure 3: Capsular polysaccharide thickness quantification. a,** Histograms of the thickness of polysaccharide layer of EcN WT and EcN Δ*kfiC* strain. The Gaussian curve were fitted to obtain mean and standard deviation of the polysaccharide layer of each strain. Inset shows SDS-PAGE gel. Alcian blue stain confirmed presence of ~180 kDa band for EcN strain. **b,** TEM image processing of polysaccharide layer. Raw images were first processed using Gaussian blur to reduce noise and further transformed into binary images for image analysis to measure the thickness of ruthenium red-stained polysaccharide layer.

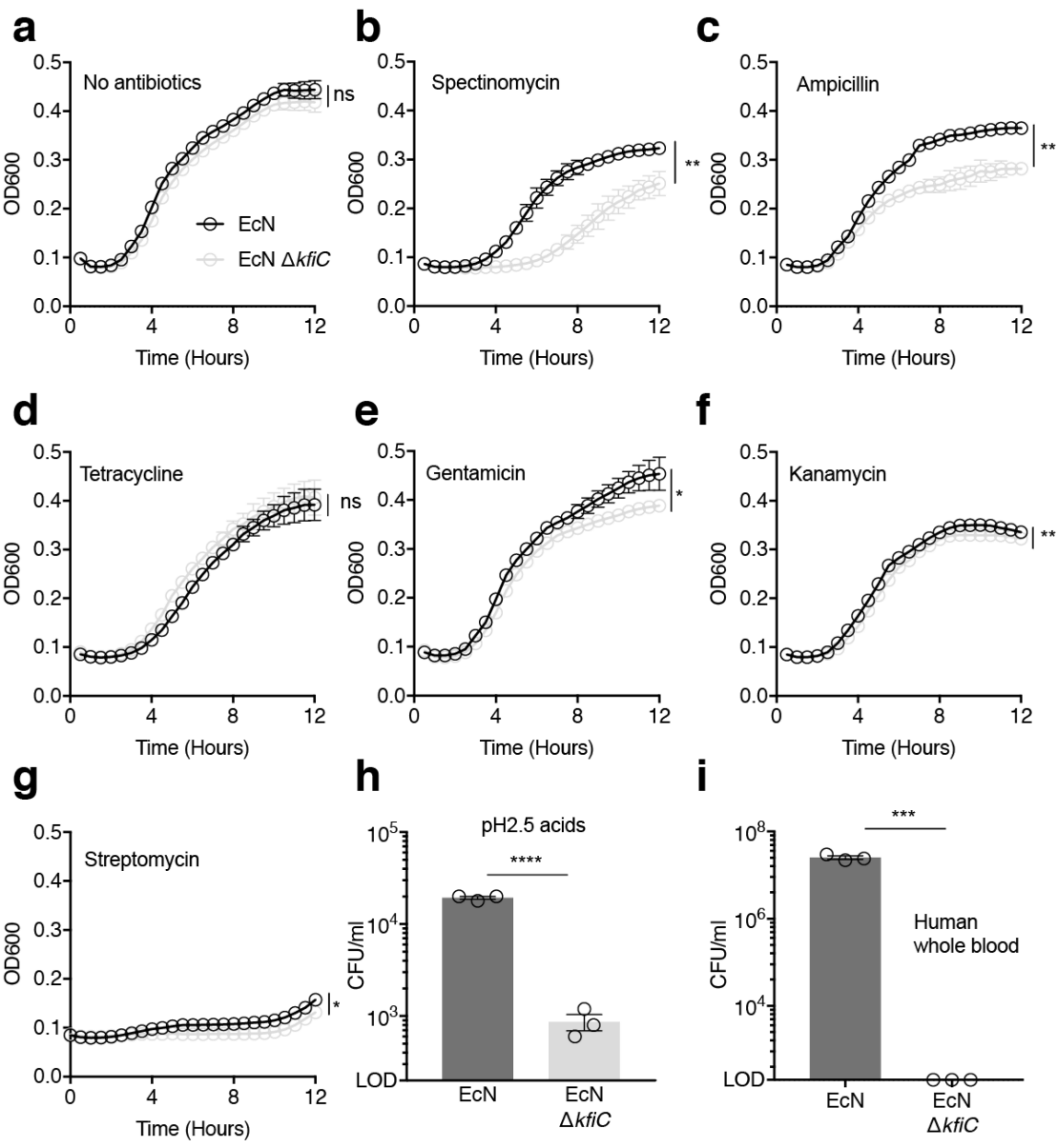


**Supplementary Figure 4: Bacterial protection against environmental threats. a–g,** Growth kinetics of EcN and EcN Δ*kfiC* in LB media containing sublethal concentration of antibiotics. (a) No antibiotics, (b) 10 µg/mL Spectinomycin, (c) 2 µg/mL Ampicillin, (d) 0.2 µg/mL Tetracycline, (e) 1 µg/mL Gentamicin, (f) 10 µg/mL Kanamycin, (g) 5 µg/mL Streptomycin (n.s. = 0.0585, **P = 0.0043, **P = 0.0042, n.s. = 0.255, *P = 0.041, **P = 0.0089, *P = 0.016 respectively. n = 3. two-way ANOVA). **h,** Bacterial survival in low pH condition. Bacteria were incubated in LB media at pH = 2.5 for 1 hour and plated on LB agar for CFU enumeration. **i,** Bacterial survival in human blood. Bacteria were inoculated in human whole blood for 0.5 hours and plated on LB agar for CFU enumeration. All error bars represent SEM.

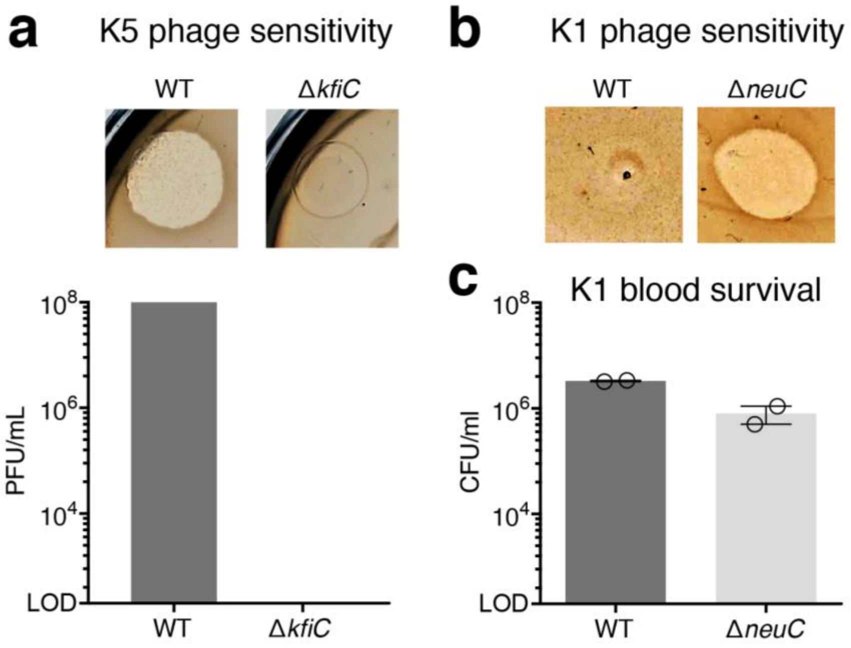


**Supplementary Figure 5: Characterization of CAP deletion in K1 and K5 strains. a,b,** Phage sensitivity of WT and KO mutant of *E. coli* (a) K5 and (b) K1 strains. K1 CAP protects against T7 phage, and K5 CAP is targeted by ΦK1-5 phage. Quantification of K5 plaque assay is shown at the bottom bar plot. **c,** K1 bacterial survival in serum. WT and Δ*neuC* K1 strains were inoculated in mouse serum for 1.5 hour and plated on LB agar for CFU enumeration. All error bars represent SEM.

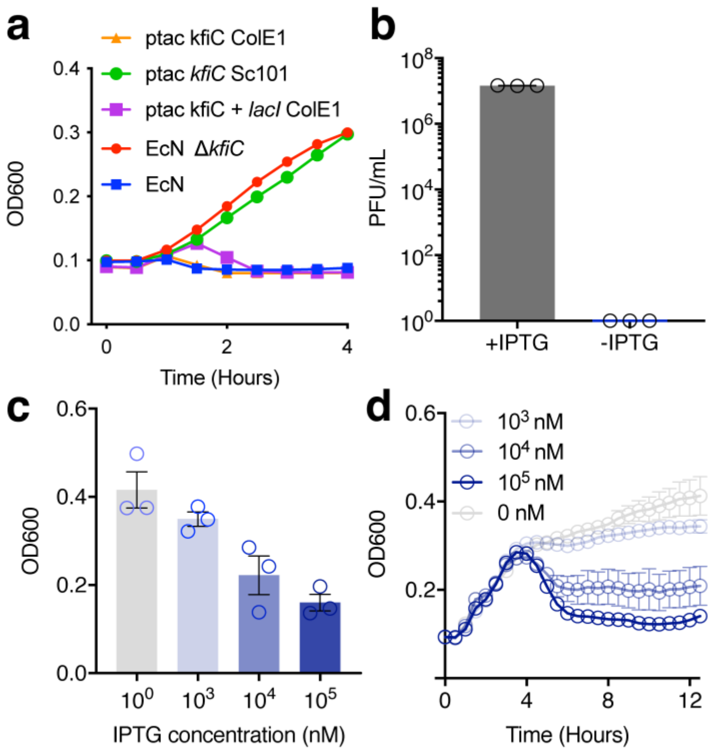


**Supplementary Figure 6: Phage sensitivity of the programmable capsular polysaccharide (iCAP) system. a,** Growth curve of EcN expressing *kfiC* gene under various copy number plasmids. Bacteria were grown in LB media containing ΦK1-5. **b,** Phage plaque assay of induced and uninduced EcN iCAP. Absence of IPTG resulted in complete immunity against ΦK1-5. IPTG induction rescued sensitivity to ΦK1-5. **c,** Co-incubation of EcN Δ*kfiC* transformed with plasmids encoding *kfiC* and *lacI* (referred to as EcN iCAP hereafter) and ΦK1-5 in varying concentrations of IPTG. Inversely proportional relationship between IPTG concentration and viability of EcN iCAP were observed. **d,** Growth curve of EcN iCAP in LB media containing ΦK1-5. iCAP was pre-induced in various IPTG concentrations. Upon inoculation, rapid bacteria lysis event was observed after 3.5 hours. Inversely proportional relationship between IPTG concentration and bacteria lysis were observed. All error bars represent SEM.

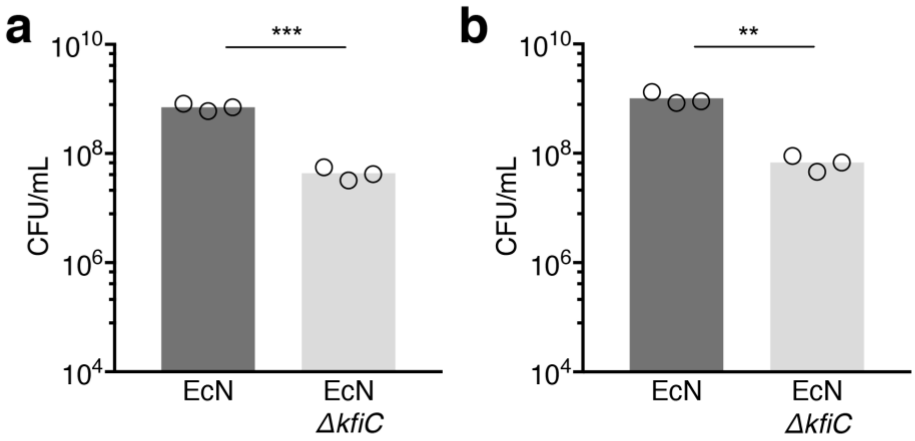


**Supplementary Figure 7: Bacterial survival in mouse whole blood. a,b,** EcN and EcN Δ*kfiC* were inoculated in mouse whole blood for (a) 1 and (b) 2 hours and plated on LB agar for CFU enumeration (***P = 0.0005, **P = 0.003. Unpaired t-test).

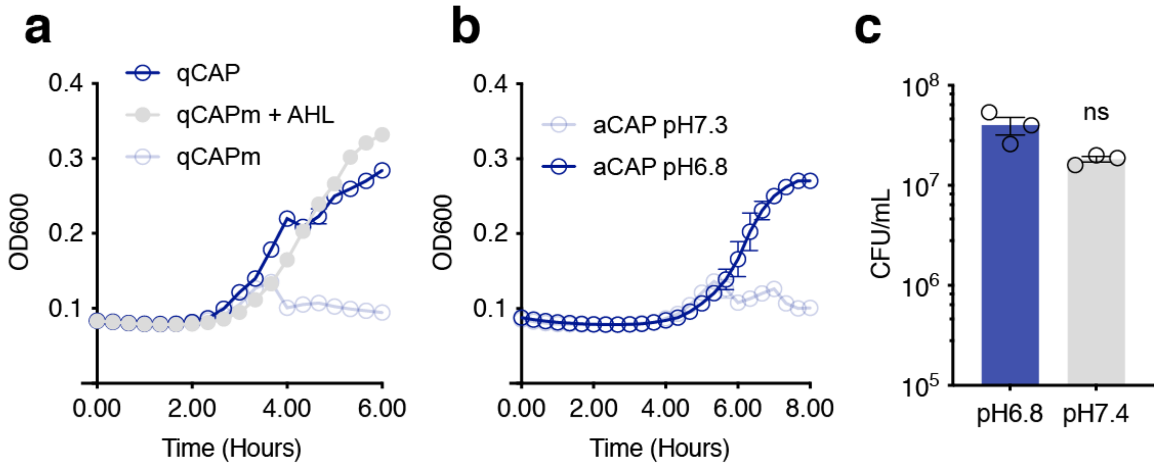


**Supplementary Figure 8: Biosensor regulates CAP-mediated survival in phage and blood. a,** EcN qCAP were grown in media containing ΦK1-5 to measure phage sensitivity. qCAP strain that was allowed to reach stationary phase before inoculating phage was able to grow in the media, suggesting that CAP is suppressed. qCAPm denotes a control strain with mutated *luxI* gene, unable to produce functional AHL to reach quorum. This strain was sensitive to ΦK1-5, suggesting CAP expression. Exogenous addition of 10nM AHL allowed for qCAPm strain, supporting that the CAP expression in this sytem is mediated by quorum sensing mechanism. **b,** EcN aCAP were grown in media containing ΦK1-5 to measure phage sensitivity at neutral (pH 7.3) or acidic (pH 6.8) condition. While the bacterial growth was suppressed at neutral condition, aCAP growth was observed in acidic media, suggesting that CAP is repressed in acidic condition. **c,** Control strain with constitutive *kfiC* expression was inoculated in neutral or acidic human whole blood. Blood pH did not affect bacterial survival without aCAP sensing gene circuit.

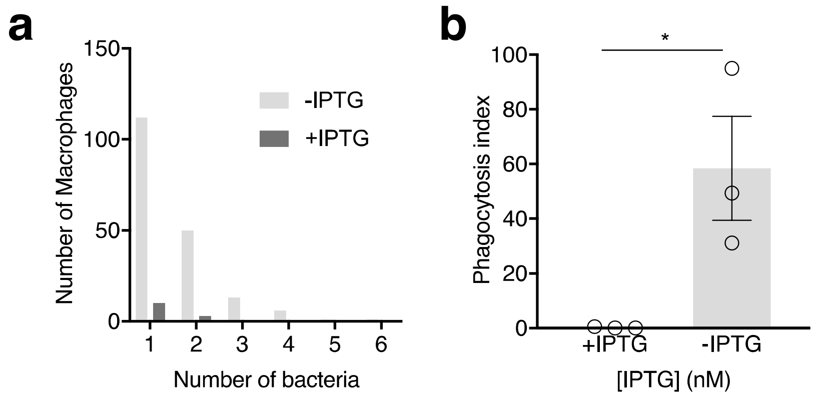


**Supplementary Figure 9: Inducible protection from phagocytosis using iCAP. a,** Histogram showing number of phagocytosed bacteria in murine bone marrow derived macrophages (BMDM). **b,** iCAP activation reduced levels of phagocytosis by BMDM (*P = 0.037, t-test). Phagocytosis index = (% BMDM containing >1 bacterium) × (mean number of bacteria per BMDM). All error bars represent SEM.

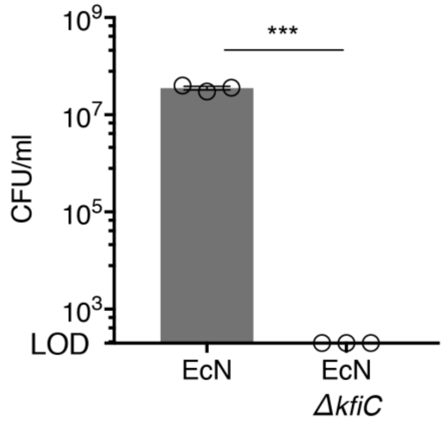


**Supplementary Figure 10: Bacterial survival in human plasma.** EcN and EcN Δ*kfiC* were inoculated in mouse whole blood for 0.5 hour and plated on LB agar for CFU enumeration. All error bars represent SEM.

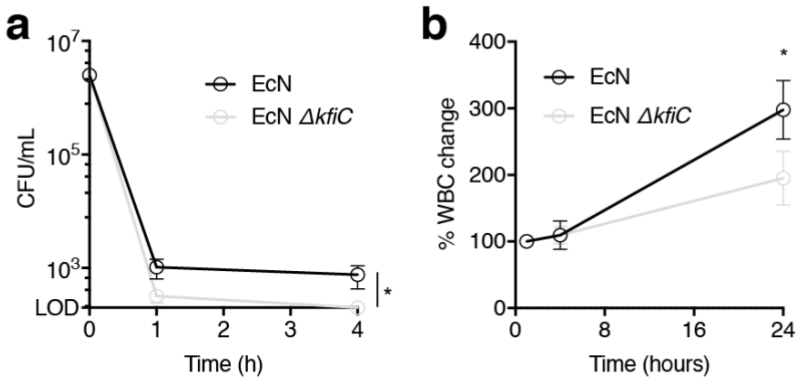


**Supplementary Figure 11: Change in white blood cell (WBC) count in blood.** EcN induced greater levels of WBC expansion compared to EcN Δ*kfiC*. Difference in WBC levels were observed after 24 hours p.i. (*P = 0.033, two-way ANOVA with Sidak's multiple comparisons test, n = 5 per group). All error bars represent SEM.

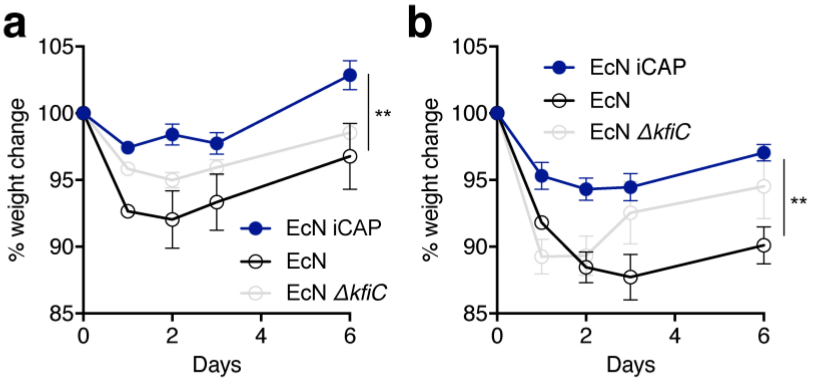


**Supplementary Figure 12: Change in animal body weight after intravenous bacterial administration at varying doses. a,b,** Bacteria were intravenously administered to BALB/c mice at (a) 5 x $10^6$ CFU and (b) 1 x $10^7$ CFU. iCAP group showed minimal drop in weight compared to EcN and EcN Δ*kfiC* groups. (**P = 0.004 and 0.001, two-way ANOVA with Turkey's multiple comparison test. n = 10, 5, 5 mice per group, respectively, for EcN iCAP, EcN, and EcN Δ*kfiC*). All error bars represent SEM.

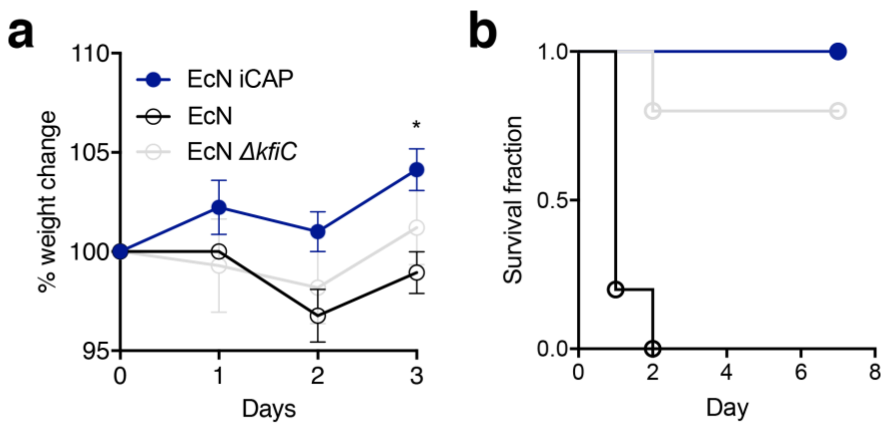


**Supplementary Figure 13: Toxicity characterization of iCAP strains in sepsis model.** **a,** $10^6$ CFU bacteria were intraperitoneally administered to BALB/c mice. iCAP group showed minimal drop in weight compared to EcN and EcN Δ*kfiC* groups. (*P = 0.0153, two-way ANOVA with Turkey's multiple comparison test. n = 5 mice per group). All error bars represent SEM. **b,** Survival curve after $10^7$ CFU bacterial administration. Animals injected with EcN iCAP all survived while EcN group all succumbed within 2 days.

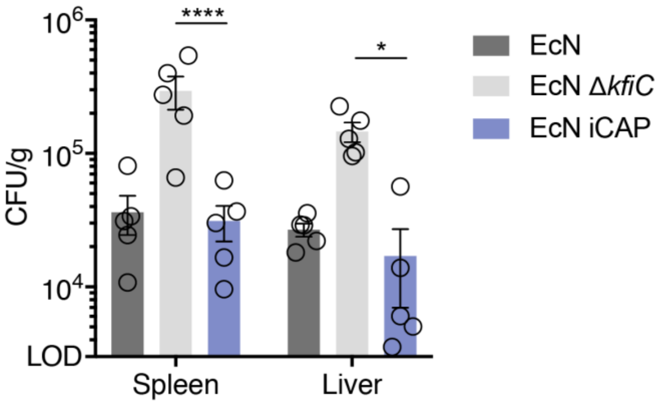


**Supplementary Figure 14: Bacterial biodistribution upon intravenous delivery *in vivo*.** BALB/c mice were intravenously administered with EcN, EcN Δ*kfiC*, or EcN iCAP. EcN iCAP was pre-induced with 10 µM IPTG. Spleen and liver were harvested after 1 day, homogenized, and spotted on LB-agar plate for CFU enumeration. Transient protection by EcN iCAP demonstrated reduced CFU in peripheral organs compared to EcN Δ*kfiC*. (****P < 0.0001, *P = 0.0452, two-way ANOVA with Turkey's multiple comparison test. LOD = 3 x $10^6$ CFU/g). All error bars represent SEM.

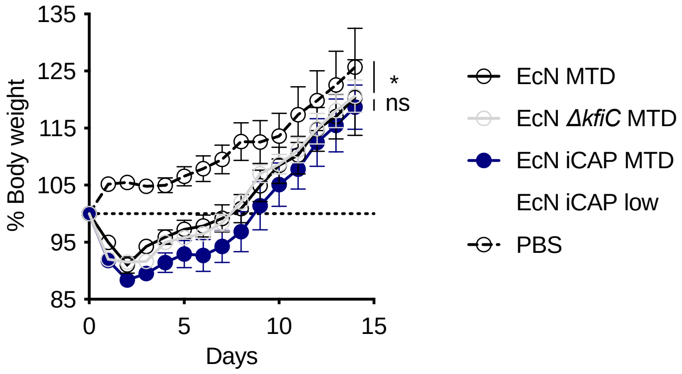


**Supplementary Figure 15: Change in animal body weight after bacterial administration at MTD.** Mice were i.v. injected with EcN MTD, EcN *ΔkfiC* MTD, EcN iCAP MTD (pre-induced with 10 µM IPTG), or EcN iCAP low (pre-induced with 10 µM IPTG) expressing antitumor theta-toxin at 5x10$^{6}$, 1x10$^{7}$, 5x10$^{7}$, or 5x10$^{6}$ CFU, respectively. All animals showed similar drop in body weight at MTD (*P = 0.03, n.s. P > 0.49; two-way ANOVA with Turkey's multiple comparison test; n > 4 mice per group). All error bars represent SEM.

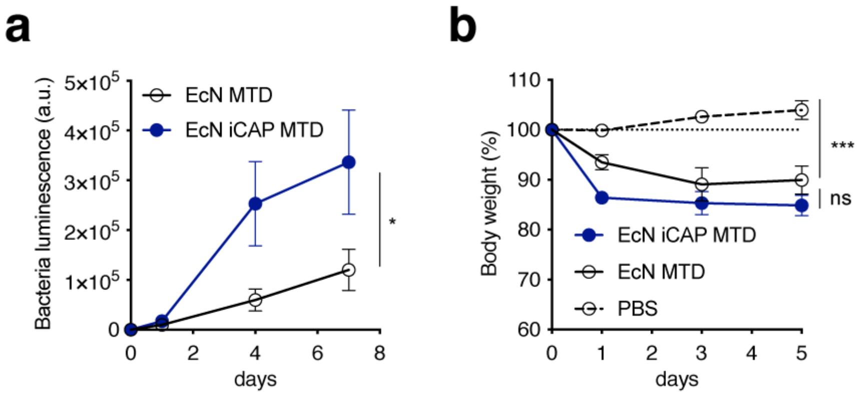


**Supplementary Figure 16:** Bacterial administration at MTD in PyMT-MMTV model. **a,** Bacterial growth trajectories in PyMT tumors after intravenous delivery *in vivo.* Each line represents average of bacterial growth trajectories in tumors quantified by bacterial luminescence over time for each bacterial strain injected. Tumors injected with EcN iCAP MTD showed higher bacterial luminescence compared to tumor injected with EcN MTD (*P = 0.044, Two-way ANOVA with Turkey's multiple comparison test; n = 15 tumors for EcN MTD and EcN iCAP MTD groups). Luminescence values are normalized to basal luminescence of individual strains. **b,** Change in animal body weight after bacterial administration at MTD. Mice were i.v. injected with EcN MTD or EcN iCAP MTD (pre-induced with 10 µM IPTG) expressing antitumor theta-toxin. Both groups showed similar drop in animal body weight (***P = 0.0002, n.s. P > 0.1; two-way ANOVA with Turkey's multiple comparison test; n > 3 mice per group). All error bars represent SEM.

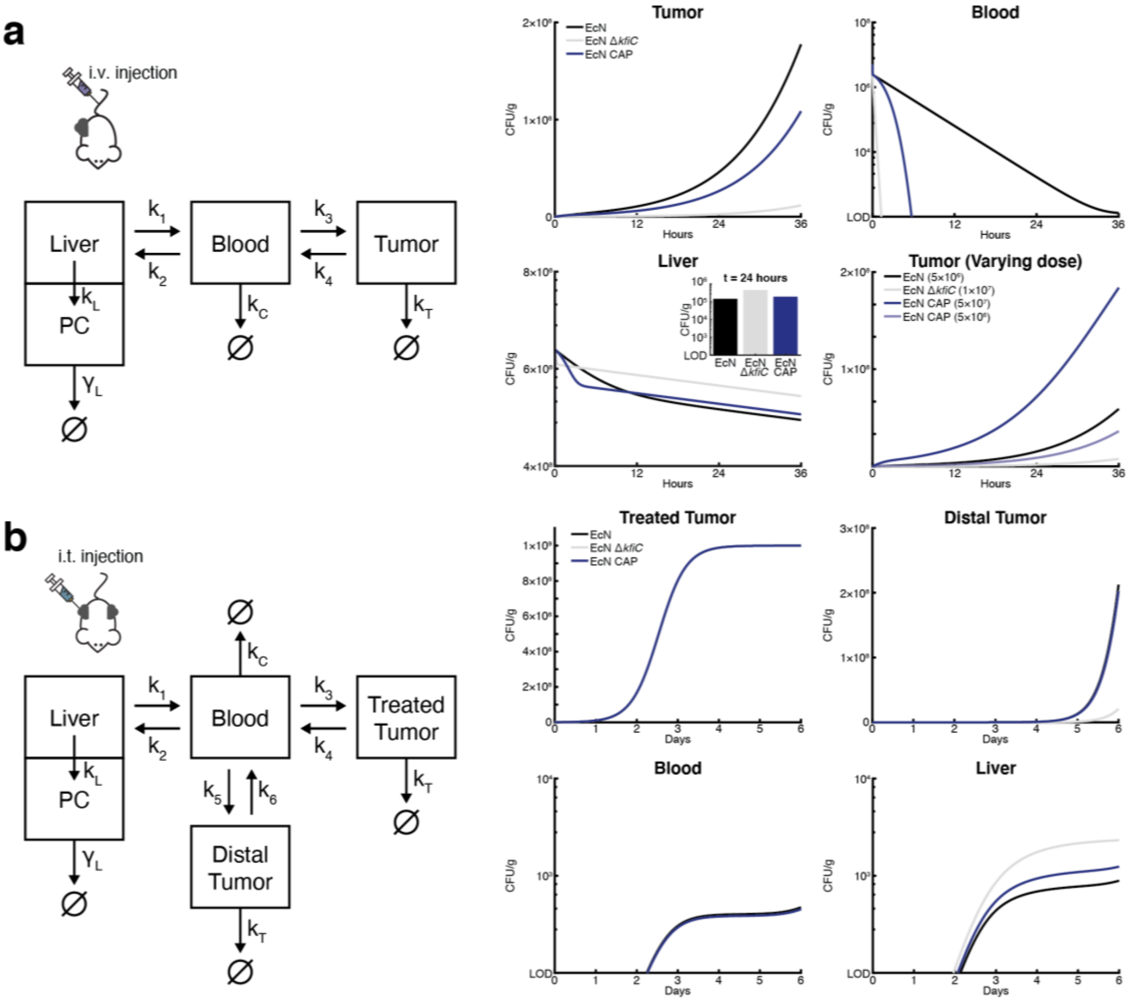


**Supplementary Figure 17. Bacterial pharmacokinetics model. a,** A 3-compartment pharmacokinetic (PK) model for delivery via i.v. injection. To simulate the i.v. injection, initial conditions are set so that the bacterial population in each compartment other than blood is equal to zero. The magnitude of initial condition in blood acts as the different injection doses. **b,** A 4-compartment PK model for delivery via i.t. injection. To simulate the i.t. injection, the initial conditions are set so that the bacterial population in each compartment other than treated tumor is equal to zero.

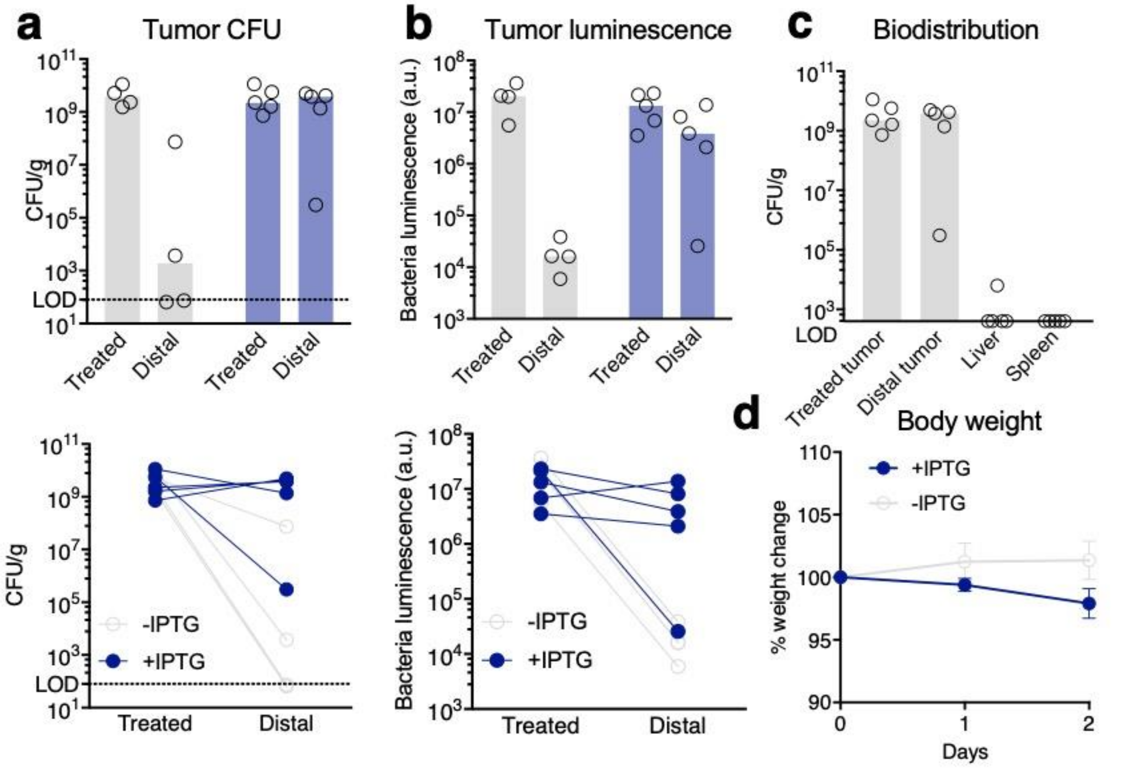


**Supplementary Figure 18: Inducible translocation of EcN iCAP in CT26 model. a,** Inducible translocation of EcN iCAP from treated tumors to distal tumors in CFU. Mice bearing subcutaneous CT26 tumors were injected intratumorally with EcN iCAP to one tumor (treated). One group was fed with water containing IPTG 1 day p.i. (+IPTG in blue) to activate iCAP *in situ*. Top graphs represent bacterial CFU in tumors. Tumors were harvested after 3 days p.i., homogenized, and spotted on LB-agar plate for CFU enumeration. Bars denotes medians. Bottom graphs represent bacterial CFU connected with lines showing individual tumor pairs. **b,** Inducible translocation of EcN iCAP from treated tumors to distal tumors quantified by bacterial luminescence. Top graphs represent bacterial luminescence in tumors, corresponding to IVIS images. Tumors were harvested after 3 days p.i. and images *ex vivo*. Bars denotes medians. Bottom graphs represent bacterial luminescence connected with lines showing individual tumor pairs. **c,** Bacterial biodistribution upon intratumoral administration and translocation *in vivo*. Tumors, spleen and liver were harvested after 3 days p.i., homogenized, and spotted on LB-agar plate for CFU enumeration. *In situ* induction of EcN iCAP demonstrated colonization of distal tumors. Bars denotes median. **d,** Change in animal body weight after intratumoral bacterial administration and induced translocation. Graphs represent % change in animal body weight p.i. All error bars represent SEM.

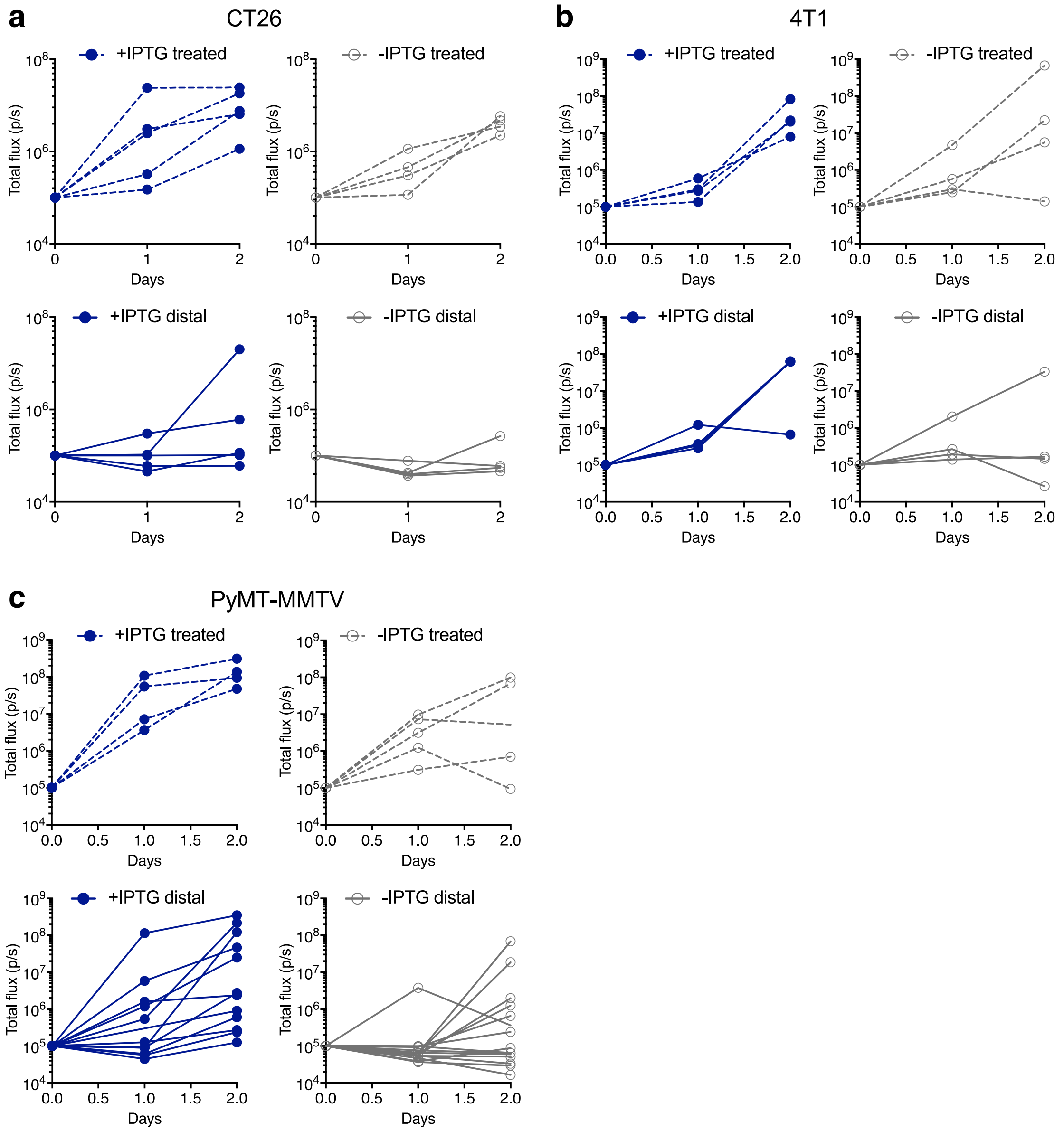


**Supplementary Figure 19: Individual bacteria growth trajectories in tumors after intratumoral administration of single tumor flank *in vivo*. a-c,** Mice bearing either (a) subcutaneous CT26, (b) orthotropic 4T1, or (c) spontaneous PyMT-MMTV tumors were injected intratumorally with EcN iCAP to one tumor (treated, dotted lines). One group was fed with water containing IPTG 1 day p.i. (+IPTG, blue lines) to activate iCAP *in situ*. Graphs represent individual bacterial growth trajectories in tumors quantified by bacterial luminescence over time, corresponding to IVIS images. Increasing level of bacterial luminescence in untreated tumors (distal, solid lines) was observed in groups induced with IPTG.

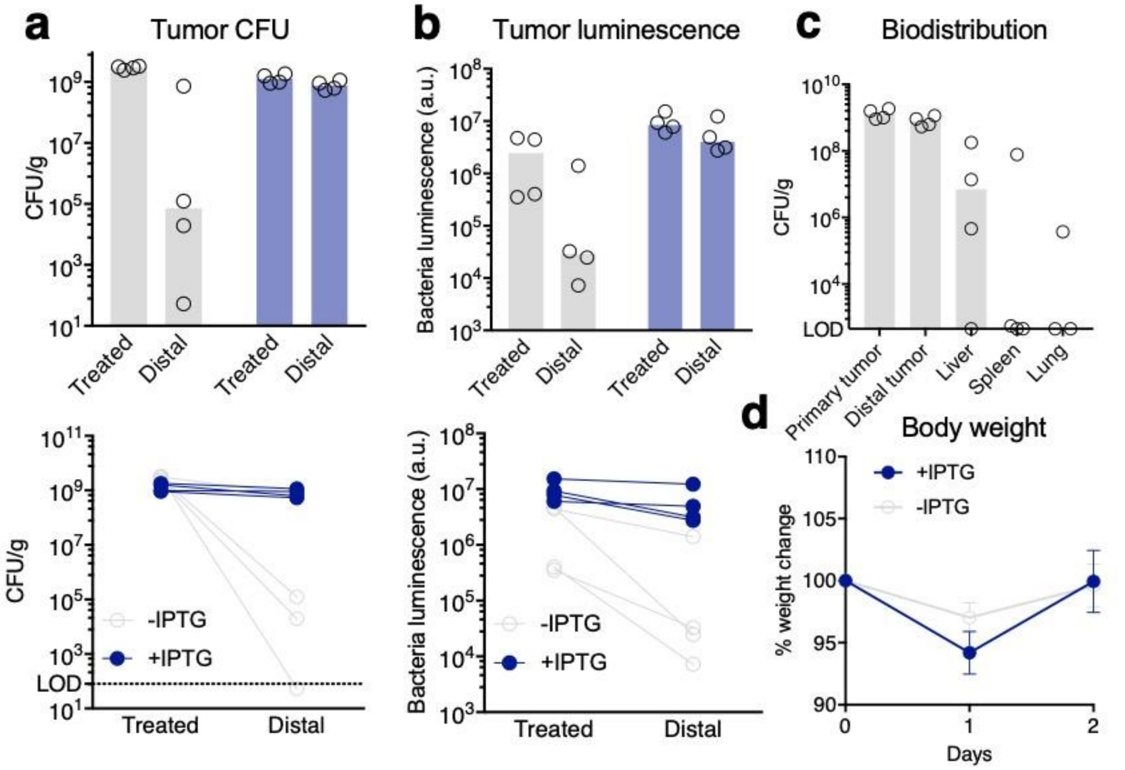


**Supplementary Figure 20: Inducible translocation of EcN iCAP in 4T1 model. a,** Inducible translocation of EcN iCAP from treated tumors to distal tumors in CFU. Mice bearing orthotropic 4T1 tumors were injected intratumorally with EcN iCAP to one tumor (treated). One group was fed with water containing IPTG 1 day p.i. (+IPTG in blue) to activate iCAP *in situ*. Top graphs represent bacterial CFU in tumors. Tumors were harvested after 3 days p.i., homogenized, and spotted on LB-agar plate for CFU enumeration. Bars denotes medians. Bottom graphs represent bacterial CFU connected with lines showing individual tumor pairs. **b,** Inducible translocation of EcN iCAP from treated tumors to distal tumors quantified by bacterial luminescence. Top graphs represent bacterial luminescence in tumors, corresponding to IVIS images. Tumors were harvested after 3 days p.i. and images *ex vivo*. Bars denotes medians. Bottom graphs represent bacterial luminescence connected with lines showing individual tumor pairs. **c,** Bacterial biodistribution upon intratumoral administration and translocation *in vivo*. Tumors, spleen and liver were harvested after 3 days p.i., homogenized, and spotted on LB-agar plate for CFU enumeration. *In situ* induction of EcN iCAP demonstrated colonization of distal tumors. Bars denotes median. **d,** Change in animal body weight after intratumoral bacterial administration and induced translocation. Graphs represent % change in animal body weight p.i. All error bars represent SEM.

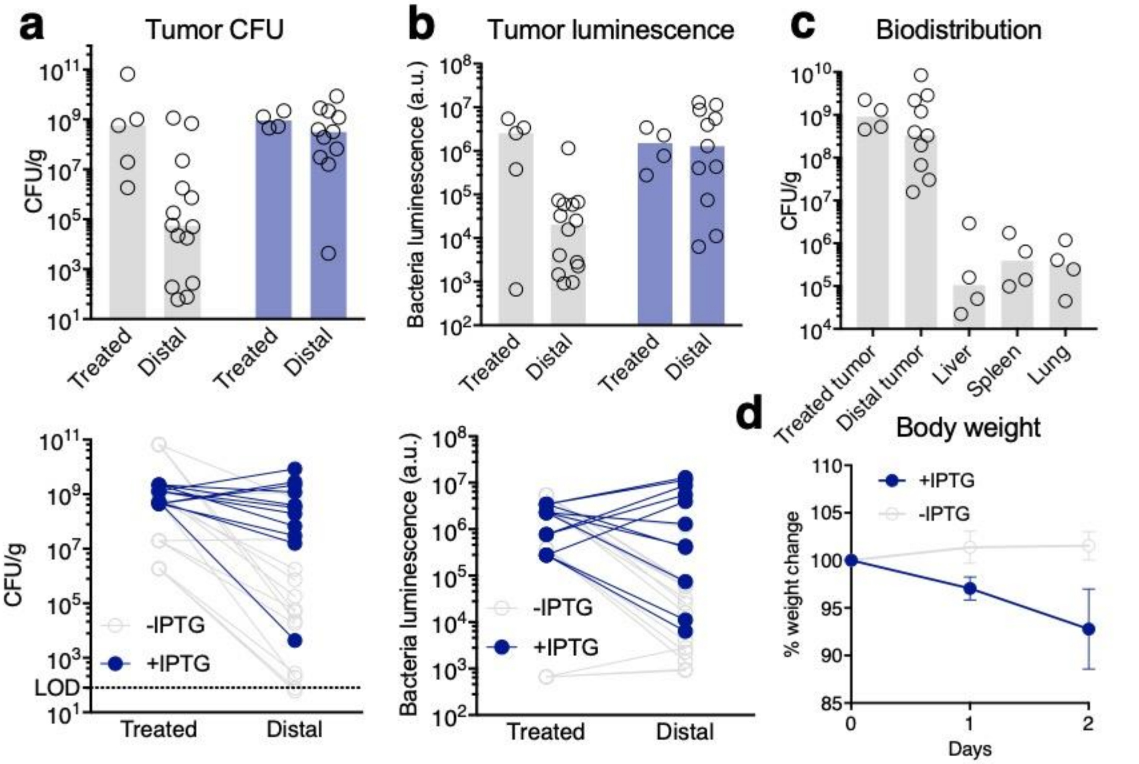


**Supplementary Figure 21: Inducible translocation of EcN iCAP in PyMT-MMTV model. a,** Inducible translocation of EcN iCAP from treated tumors to distal tumors in CFU. Mice bearing spontaneous PyMT-MMTV tumors were injected intratumorally with EcN iCAP to one tumor (treated). One group was fed with water containing IPTG 1 day p.i. (+IPTG in blue) to activate iCAP *in situ*. Top graphs represent bacterial CFU in tumors. Tumors were harvested after 3 days p.i., homogenized, and spotted on LB-agar plate for CFU enumeration. Bars denotes medians. Bottom graphs represent bacterial CFU connected with lines showing individual tumor pairs. **b,** Inducible translocation of EcN iCAP from treated tumors to distal tumors quantified by bacterial luminescence. Top graphs represent bacterial luminescence in tumors, corresponding to IVIS images. Tumors were harvested after 3 days p.i. and images *ex vivo*. Bars denotes medians. Bottom graphs represent bacterial luminescence connected with lines showing individual tumor pairs. **c,** Bacterial biodistribution upon intratumoral administration and translocation *in vivo*. Tumors, spleen and liver were harvested after 3 days p.i., homogenized, and spotted on LB-agar plate for CFU enumeration. *In situ* induction of EcN iCAP demonstrated colonization of distal tumors. Bars denotes median. **d,** Change in animal body weight after intratumoral bacterial administration and induced translocation. Graphs represent % change in animal body weight p.i. All error bars represent SEM.

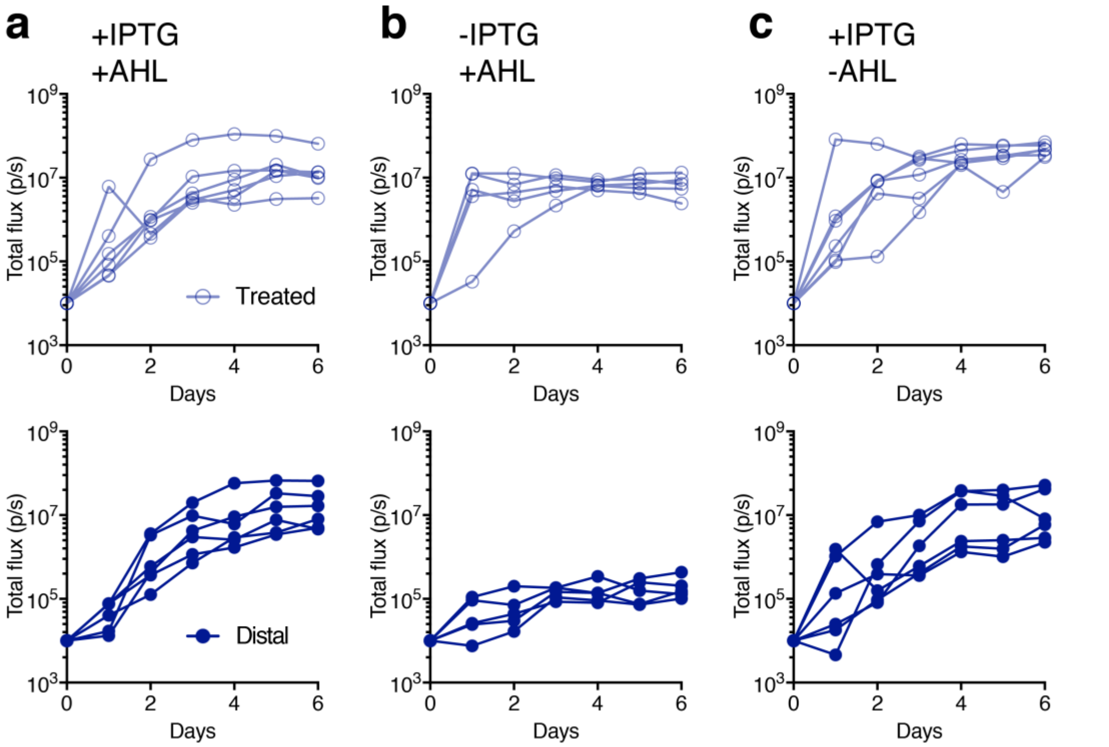


**Supplementary Figure 22: Individual growth trajectories of therapeutic bacteria in tumors after intratumoral administration of single tumor flank *in vivo*. a-c,** Mice bearing subcutaneous CT26 tumors were injected intratumorally with EcN iCAP engineered to produce TT when induced with AHL to one tumor. (a) One group was fed with water containing IPTG 1 day p.i. to activate iCAP *in situ*, and subcutaneously injected with AHL to induce TT expression (+IPTG +AHL). (b) One group only received AHL (-IPTG +AHL). (c) One group only received IPTG (+IPTG -AHL). Graphs represent individual bacterial growth trajectories in tumors quantified by bacterial luminescence over time, corresponding to IVIS images. Increasing level of bacterial luminescence in untreated tumors (distal, solid lines) was observed in groups induced with IPTG.

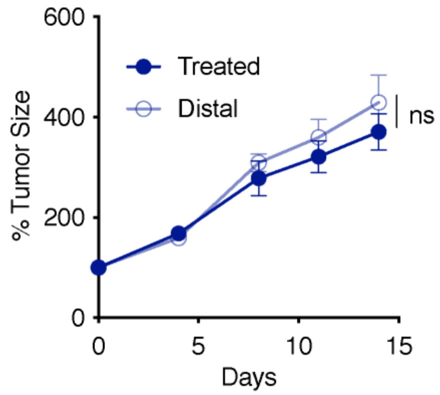


**Supplementary Figure 23: Treated and distal CT26 tumors from EcN iCAP control group measured by relative tumor growth over time.** Bacteria were injected into a single treated tumor. The translocation was controlled by IPTG water. After 3 days of initial injection, AHL were administered to induce therapeutic expression. iCAP control does not contain theta-toxin gene. (n.s. $P = 0.92$, two-way ANOVA with Bonferroni posttest, $n = 6$ for both treated and distal tumors). All error bars represent standard error of mean (SEM).

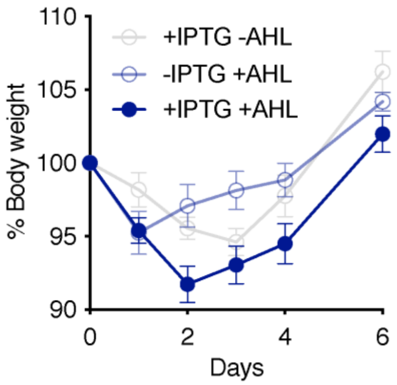


**Supplementary Figure 24: Change in animal body weight after intratumoral therapeutic bacterial administration and induced translocation.** Mice bearing subcutaneous CT26 tumors were injected intratumorally with EcN iCAP engineered to produce TT when induced with AHL to one tumor. One group was fed with water containing IPTG 1 day p.i. (+IPTG -AHL) to activate iCAP *in situ*. One group was subcutaneously injected with AHL to induce TT expression (-IPTG +AHL). One group received both IPTG and AHL (+IPTG +AHL). Graphs represent % change in animal body weight p.i. All error bars represent SEM.

| Identifier | Bacterial Strains | Plasmids (ORI/promoter) | Relevant Features | Figure |
|---|---|---|---|---|
| EcN | *E. coli* Nissle 1917 | N/A | N/A | 2, 4, 5, S1-4, S7, S10-15 |
| K1 | *E. coli* K1 strain | N/A | N/A | S1, S5 |
| K5 | *E. coli* K5 strain | N/A | N/A | S5 |
| MG1655 | *E. coli* MG1655 strain | N/A | N/A | S1 |
| VNP20009 | *S. typhimurium* VNP20009 | N/A | N/A | S1 |
| EcN KD | *E. coli* Nissle 1917 | ColE1/pPR | sRNA with MicC scaffold binding to gene of interest | 2, S2 |
| EcN KO | *E. coli* Nissle 1917 | N/A | Genomic deletion of *kfi* genes | S2 |
| EcN Δ*kfiC* | *E. coli* Nissle 1917 Δ*kfiC* | N/A | Genomic deletion of *kfiC* gene | 2, 4, 5, S2-4, S7, S10-15 |
| EcN iCAP | *E. coli* Nissle 1917 Δ*kfiC* | Sc101/ptac | *kfiC* gene expressed under ptac promoter | 3-6, S6, S9, S12-15, S17-20 |
| EcN qCAP | *E. coli* Nissle 1917 Δ*kfiC* | ColE1/ptac<br><br>P15A/pluxI | *kfiC* gene expressed under ptac promoter<br><br>*luxI* and *lacI* gene expressed under pluxI promoter | 4, S8 |
| EcN aCAP | *E. coli* Nissle 1917 Δ*kfiC* | ColE1/ptac<br><br>P15A/pCadC | *kfiC* gene expressed under ptac promoter<br><br>*lacI* gene expressed under pCadC promoter | 4, S8 |
| EcN iCAP TT | *E. coli* Nissle 1917 Δ*kfiC* | Sc101/ptac<br><br>ColE1/pluxI | *kfiC* gene expressed under ptac promoter<br><br>*theta* gene expressed under pluxI promoter | 5, 6, S15, S21-23 |

**Supplementary Table 1.** Main bacterial strains and plasmids used in this study.

| Parameter | Definition | Value |
| --- | --- | --- |
| g | Bacterial growth rate | 13 |
| K | Bacteria carrying capacity | 0.05 |
| $k_1$ | Transfer constant from blood to liver | 1000 |
| $k_2$ | Transfer constant from liver to blood | 1000 |
| $k_3$ | Transfer constant from blood to (treated) tumor | 10 |
| $k_4$ | Transfer constant from (treated) tumor to blood | 0.00001 |
| $k_5$ | Transfer constant from blood to distal tumor | 10 |
| $k_6$ | Transfer constant from distal tumor to blood | 0.00001 |
| $\gamma_L$ | Rate of phagocytic lysis in liver | 1 |
| $\sigma_C$ | Rate of complement-mediated lysis | 15 (EcN/+CAS)<br>50 (EcN/-CAS) |
| $C_{50}$ | Maximum concentration of complements | 1 |
| $\sigma_T$ | Phagocytic elimination rate in the tumor | 10 |
| $\sigma_L$ | Phagocytic capturing rate in the liver | 0.01 (EcN/+CAS)<br>5 (EcN/-CAS) |
| $M_T$ | Maximum bacterial clearance in tumor | 1 |
| $M_L$ | Maximum bacterial clearance in liver | 1 |
| $x$ | Non linearity in phagocytic capturing function | 2 |
| $\mu$ | Logistic growth rate of switch parameters | 10 ($\sigma_C$, IV)<br>20 ($\sigma_L$, IV)<br>10 ($\sigma_C$, IT)<br>20 ($\sigma_L$, IT) |

**Supplementary Table 2.** Parameters used for the simulations.

#### Supplementary Note on Computational Model Derivation

To simulate the probiotic biodistribution resulting from the programmable probiotic encapsulation system, we used pharmacokinetic (PK) compartmental modeling. Due to the scope of the paper and its modeling requirements, we examined biodistribution to only the most essential compartments. We adopted relatively simple methods used to model nanoparticle delivery [1,2] to determine how the compartments are constructed. Thus, our model contains a single blood compartment as opposed to distinct arterial and venous compartments, and the liver compartment has an additional subcompartment for phagocytosis-captured bacteria. We assume each major compartment to have the same bacterial carrying capacity (*i.e.*, same volume and maximum concentration of bacteria).

##### A. Three-Compartment PK Model for Intravenous Delivery

We applied a three-compartment PK model for delivery via intravenous (i.v.) injection (Supplementary Fig. 17a). As mentioned above, the compartments include blood, liver, and tumor. Tumor was selected as one of the three compartments because it is the therapeutic target. Blood and liver were selected because they contain the main modalities for host immune clearance of injected bacteria.

Movements of bacteria between the main compartments (blood, liver, and tumor) are determined by transfer constants $k_{1\text{–}4}$ and the amount of bacteria available in each compartment. The blood compartment is the facilitator of bacteria distribution between the other periphery compartments. Transfer between the blood and liver compartments is governed by $k_1$ and $k_2$, and transfer between the blood and tumor compartments is governed by $k_3$ and $k_4$. The various functions of the host immune system are governed by $k_C$, $k_L$, $k_T$ and $\gamma_L$. $k_C$ is the rate constant of bacterial elimination in the blood by way of complement-mediated lysis, $k_L$ is the rate constant of phagocytic "capturing" in the liver, and $k_T$ is the rate constant of phagocytic elimination in the tumor. Phagocytic capturing rate constant, $k_L$, is a one-way transfer constant that moves bacteria from the main liver compartment into the phagocytic subcompartment. Once bacteria have been transfered into the phagocytic subcompartment, it cannot be transfered back into the main compartment. The captured bacteria is eliminated at rate governed by $\gamma_L$.

As is customary in PK modeling, our model consists of a system of differential equations where relevant terms are multiplied by the bacterial population in source compartments then added together. Overall bacterial growth dynamics is modeled using logistic growth. Each compartment is assumed to have the same bacterial growth rate *g* and carrying capacity *K*. To simulate the i.v. injection itself, initial conditions are set so that the bacterial population in each compartment other than in the blood is equal to zero. The magnitude of the positive initial condition acts as the injection dosage. The full set of equations can be found below.

$$\frac{dB}{dt} = gB + k_2L + k_4T - (k_1 + k_3 + k_C)B \tag{1}$$

$$\frac{dL}{dt} = gL + k_1B - (k_2 + k_L)L \tag{2}$$

$$\frac{dT}{dt} = [gT + k_3B - (k_4 + k_T)T](1 - \frac{T}{K}) \tag{3}$$

$$\frac{dPC_L}{dt} = L - \gamma_L PC_L \tag{4}$$

$$k_C = \frac{\sigma_C}{C_{50} + B} \tag{5}$$

$$k_L = \frac{\sigma_L}{M_L L^x + 1} \tag{6}$$

$$k_T = \frac{\sigma_T}{M_T T^x + 1} \tag{7}$$

**B. Host Immune Clearance**

Bacteria in our model is eliminated by two main host immune modalities: phagocytosis in the liver and complement-mediated lysis in the blood. To model phagocytosis, we included a subcompartment that contains the population of captured bacteria still alive in phagosomes. Bacteria that enter the subcompartment cannot escape and are eventually lysed at a constant rate $\gamma_L$. The phagocytic capturing term $k_L$ is not constant but rather follows equation 6. Also not constant, the rate of phagocytic elimination from the tumor $k_T$ follows equation 7. These equations were derived by simplifying equations used on the development of a model for three-stage immune response [3]. In [3], equation 8 is used to depict the interaction between pneumococcal population and resident alveolar macrophages. This interaction is similar to the bacteria-immune response relationship we are attempting to model. The derivation for the rates of phagocytic capturing and elimination in our model is as follows:

$$\gamma_{MA}\, f(B, M_A^*) = \gamma_{MA} \times \frac{n^x M_A^*}{B^x + n^x M_A^*} \tag{8}$$

$$= \frac{\gamma_{MA}}{M_B B^x + 1} \; where \; M_B = \frac{1}{n^x M_A^*}$$

Redefine constants:

- $\sigma_i = \gamma_{MA}$
- $k_i = \gamma_{MA}\, f(i, M_i^*)$

Let *i* = organ and assume *n*, *x* constant for each organ but $\gamma_{MA}$ variable.

$$k_i = \frac{\sigma_i}{M_i i^x + 1} \tag{9}$$

The parameter definitions for equation 8 can be found in [3], and the parameter definitions for equation 9 can be found in *Parameter Selection*.

In our model, complement-mediated lysis in the blood is regulated by a single term that depends on multiple parameter constants and variable bacteria population in the blood compartment. Current methods to model complement-mediated lysis entails modeling the dynamics for each protein involved in the complement cascade and in the formation of the MAC [4,5,6]. These methods proved to be too complex for the scope of our project. Thus, we adapted the equation component describing bacterial effect on cytokine expression in

[7] to describe bacterial elimination via complement-mediated lysis in our model. The cytokines in [7] characterize an indirect host immune response. In addition, cytokines play a critical role in initiating the complement cascade [9]. The adapted equation can be found in equation 6 and corresponding parameter definitions can be found in *Parameter Selection*.

**C. Four-Compartment PK Model for Intratumoral Delivery**

To model bacterial delivery via intratumoral injection (i.t.), we applied a four-compartment PK model (Supplementary Fig. 17b). The same principles and equation structures used in the three-compartment iv. model are also used in this four-compartment model. However, the tumor compartment is split into two main compartments: treated tumor and distal tumor. This split is necessary to examine tumor trafficking observed in the experimental data. To simulate the i.t. injection itself, initial conditions are set so that the bacterial population in each compartment other than in the treated tumor is equal to zero. The full set of equations for the four-compartment model is as the following:

$$\frac{dB}{dt} = gB + k_2L + k_4T - (k_1 + k_3 + k_C)B \tag{10}$$

$$\frac{dL}{dt} = gL + k_1B - (k_2 + k_L)L \tag{11}$$

$$\frac{dT_p}{dt} = [gT_p + k_3B - (k_4 + k_{T_p})T_p](1 - \frac{T_p}{K}) \tag{12}$$

$$\frac{dT_d}{dt} = [gT_d + k_5B - (k_6 + k_{T_d})T_d](1 - \frac{T_d}{K}) \tag{13}$$

$$\frac{dPC_L}{dt} = k_LL - \gamma_L PC_L \tag{14}$$

$$k_C = \frac{\sigma_C}{C_{50} + B} \tag{15}$$

$$k_L = \frac{\sigma_L}{M_L L^x + 1} \tag{16}$$

$$k_{T_p} = \frac{\sigma_T}{M_T {T_p}^x + 1} \tag{17}$$

$$k_{T_d} = \frac{\sigma_T}{M_T {T_d}^x + 1} \tag{18}$$

**D. Switch Kinetics**

In order to model the EcN inducible capsular polysaccharides (iCAP) switch kinetics between the EcN and the EcN Δ*kfiC* host immune parameter values, we apply logistic growth where each parameter $\sigma_C$ and $\sigma_L$ share different growth rates $\mu$. Parameter switch growth rates $\mu$ were adjusted so that the parameters $\sigma_L$ and $\sigma_C$ would plateau at approximately the same time. Using the logistic growth equation, we are able to precisely control the EcN iCAP parameter values associated with EcN and EcN Δ*kfiC* as well as the speed of the switch kinetics. The EcN iCAP parameters switch from those of EcN to EcN

Δ*kfiC* for three-compartment i.v. model whereas the parameters switch from those of EcN Δ*kfiC* to EcN for the four-compartment i.t. model.

**E. Parameter Selection**

To select parameters, we initially followed the relative values found in [8]. Therefore, transfer of bacteria between the blood and liver compartments is much greater than between the blood and tumor compartments. Afterwards, we manipulated the remaining parameters so that the model would produce simulations that resemble trends found in the experimental data. To replicate the CFU levels found in the experimental data, the simulation values are scaled up by a constant defined by the ratio of the assumed experimental carrying capacity ($1*10^9$) to the simulation carrying capacity (0.05). The complete list of parameters and their corresponding definitions can be found in Supplementary Table 2.

**F. Limitations**

The computational model is primarily limited by its implementation as a system of differential equations. Solutions to differential equations are unable to reach absolute zero. As a result, tumor biodistribution at all capsule settings in both the i.v. and i.t. models always reach carrying capacity. However, the experimental results show that tumor trafficking can be suppressed in i.t. models when EcN iCAP is not induced (Fig. 6c).
